# Decoding neuronal wiring by joint inference of cell identity and synaptic connectivity

**DOI:** 10.1101/2025.03.04.640006

**Authors:** Himanshu Pawankumar Gupta, Anthony W. Azevedo, Yu-C-hieh David Chen, Kristi Xing, Peter A Sims, Erdem Varol, Richard S. Mann

**Affiliations:** Zuckerman Mind Brain Behavior Institute, Columbia University, New York, NY, USA; Department of Neurobiology and Biophysics, University of Washington, WA, USA; Department of Biology, New York University, New York, NY, USA; Barnard College, Columbia University, New York, NY, USA; Department of Systems Biology, Columbia University Irving Medical Center, New York, NY, USA; Sulzberger Columbia Genome Center, Columbia University Irving Medical Center, New York, NY, USA; Department of Computer Science & Engineering at Tandon School of Engineering, New York University, New York, NY, USA; Neuroscience Institute, Langone Medical Center, New York University, New York, NY, USA; Department of Biochemistry and Molecular Biophysics, Columbia University, New York, NY, USA

## Abstract

Animal behaviors are executed by motor neurons (MNs), which receive information from complex pre-motor neuron (preMN) circuits and output commands to muscles. How motor circuits are established during development remains an important unsolved problem in neuroscience. Here we focus on the development of the motor circuits that control the movements of the adult legs in *Drosophila melanogaster*. After generating single-cell RNA sequencing (scRNAseq) datasets for leg MNs at multiple time points, we describe the time course of gene expression for multiple gene families. This analysis reveals that transcription factors (TFs) and cell adhesion molecules (CAMs) appear to drive the molecular diversity between individual MNs. In parallel, we introduce ConnectionMiner, a novel computational tool that integrates scRNAseq data with electron microscopy-derived connectomes. ConnectionMiner probabilistically refines ambiguous cell type annotations by leveraging neural wiring patterns, and, in turn, it identifies combinatorial gene expression signatures that correlate with synaptic connectivity strength. Applied to the Drosophila leg motor system, ConnectionMiner yields a comprehensive transcriptional annotation of both MNs and preMNs and uncovers candidate effector gene combinations that likely orchestrate the assembly of neural circuits from preMNs to MNs and ultimately to muscles.

## Introduction

A functional neural circuit that is capable of executing complex behaviors requires the establishment of a vast number of connections between many thousands of neurons. For example, the ∼69 motor neurons (MNs) that target one of 18 muscles in each leg of the fruit fly, *Drosophila melanogaster*, receive input from >1500 pre-motor neurons (preMN) via >200,000 synapses (Cheong et al., 2024; Lesser et al., 2024). Further, each preMN is connected to 100s of other interneurons within the fly’s central nervous system (CNS). How this daunting complexity of neural connections, generally referred to as the neuronal wiring problem, is established during development is an important and unsolved problem in neuroscience.

Large-scale single-cell RNA sequencing (scRNAseq) has been a powerful tool for describing neuronal diversity and provides necessary information for tackling the wiring problem because it can reveal the repertoire of molecules that are potentially expressed in each neuron, including genes that are required for generating synapses between neurons. However, linking patterns of gene expression with neuronal morphology, connectivity, and physiology is challenging. Not surprisingly, transcription factors (TFs) are the major drivers of neuronal diversity as they regulate all aspects of neural identity, ranging from genes encoding cell surface proteins (Li et al., 2017; Kurmangaliyev et al., 2019) to genes required for neurotransmitter synthesis (Hobert & Kratsios, 2019; Liu et al., 2018; Morey et al., 2008). Notably, unique combinations of TFs are required to establish the morphological identity of each neuron (Allan & Thor, 2015; Enriquez et al., 2015; Hobert, 2016). Further, recent work suggests that distinct sets of TFs may be used for specifying different aspects of neuronal identity, such as dendrite and axon morphologies (Dombrovski et al., 2025). However, for most aspects of neuron identity, the relevant target genes of these TFs are not known.

Complementing scRNAseq datasets, connectomics analysis of reconstructed 3D electron microscopy (EM) volumes provides unprecedented and detailed maps of neuronal connectivity (Azevedo et al., 2024; Takemura et al., 2023; Winding et al., 2023; Zheng et al., 2018). Complete EM connectomes are now available for the *C. elegans* nervous system (Cook et al., 2019), the *Drosophila* larval CNS (Ohyama et al., 2015; Winding et al., 2023), the adult *Drosophila* brain (Scheffer et al., 2020; Xu et al., 2020; Zheng et al., 2018), and two adult *Drosophila* ventral nerve cords (VNCs) from a male and a female (Azevedo et al., 2024; Takemura et al., 2023). However, while they reveal every synapse, on their own connectomes do not provide information on the mechanisms or molecules involved in neuronal circuit assembly. Tools that interrogate scRNAseq transcriptomes and integrate the results with connectome datasets have the potential to provide new insights into the wiring problem.

The adult *Drosophila* neuromuscular system is a powerful system for studying the interplay between molecular mechanisms and neural connectivity. Each MN has a stereotyped dendritic architecture and targets specific muscle fibers, creating a myotopic map in the CNS (Baek & Mann, 2009; Brierley et al., 2012; Enriquez et al., 2015). The coordinated activation of multiple MNs is responsible for executing a variety of complex behaviors including walking (Bidaye et al., 2014; Tuthill & Wilson, 2016), grooming (Seeds et al., 2014), escape (Card & Dickinson, 2008), flight (Dickinson & Muijres, 2016; Namiki et al., 2022), courtship (Clyne & Miesenböck, 2008), and copulation (Crickmore & Vosshall, 2013; Pavlou et al., 2016). Importantly, for each behavior, the activation of the correct set of MNs depends on the circuit of preMNs that directly synapse onto MNs. Thus, the characterization of the developmental processes and molecules that establish the stereotyped connectivity between preMNs, MNs and muscles has the potential to profoundly contribute to our understanding of how *Drosophila,* as well as other animals, establishes nervous systems that can execute a diverse set of complex behaviors.

Here, we focus on the neuromuscular system that controls the movements of the adult legs in *Drosophila*. There are about 69 MNs that target 18 muscles in each leg. Of these MNs, 29 are generated by a single stem cell lineage called LinA (a.k.a. Lin15B) (Figure 1A-D and S1). Based on single-cell clonal analysis, the birth order, dendritic and axon morphology, and muscle targeting are known for all 29 MNs that are generated from LinA (Figure S1; (Baek & Mann, 2009)). More recently, EM connectomes have revealed a preliminary map of all preMNs (Cheong et al., 2024; Lesser et al., 2024). For the 29 LinA MNs, there are 701 preMNs (i.e., local neurons) that are derived from 22 neuroblast hemilineages (Figure 1F). Here, we describe the single-cell transcriptomes for the LinA MN progeny at four developmental time points. Together with the wiring diagram that exists for these MNs – from preMNs to MNs to muscles – these data provide an opportunity to integrate connectome and transcriptome datasets. To this end, we developed a novel computational tool called ConnectionMiner, which has allowed us to fully resolve our scRNAseq data to define the transcriptomes of all 29 MNs at all four time points. In addition, we also applied ConnectionMiner to resolve the preMNs present in a previously published scRNAseq dataset (Allen et al., 2020).Together, this analysis outputs synaptic partner proteins that are predicted to establish connections between preMNs and MNs.

**Figure 1:**
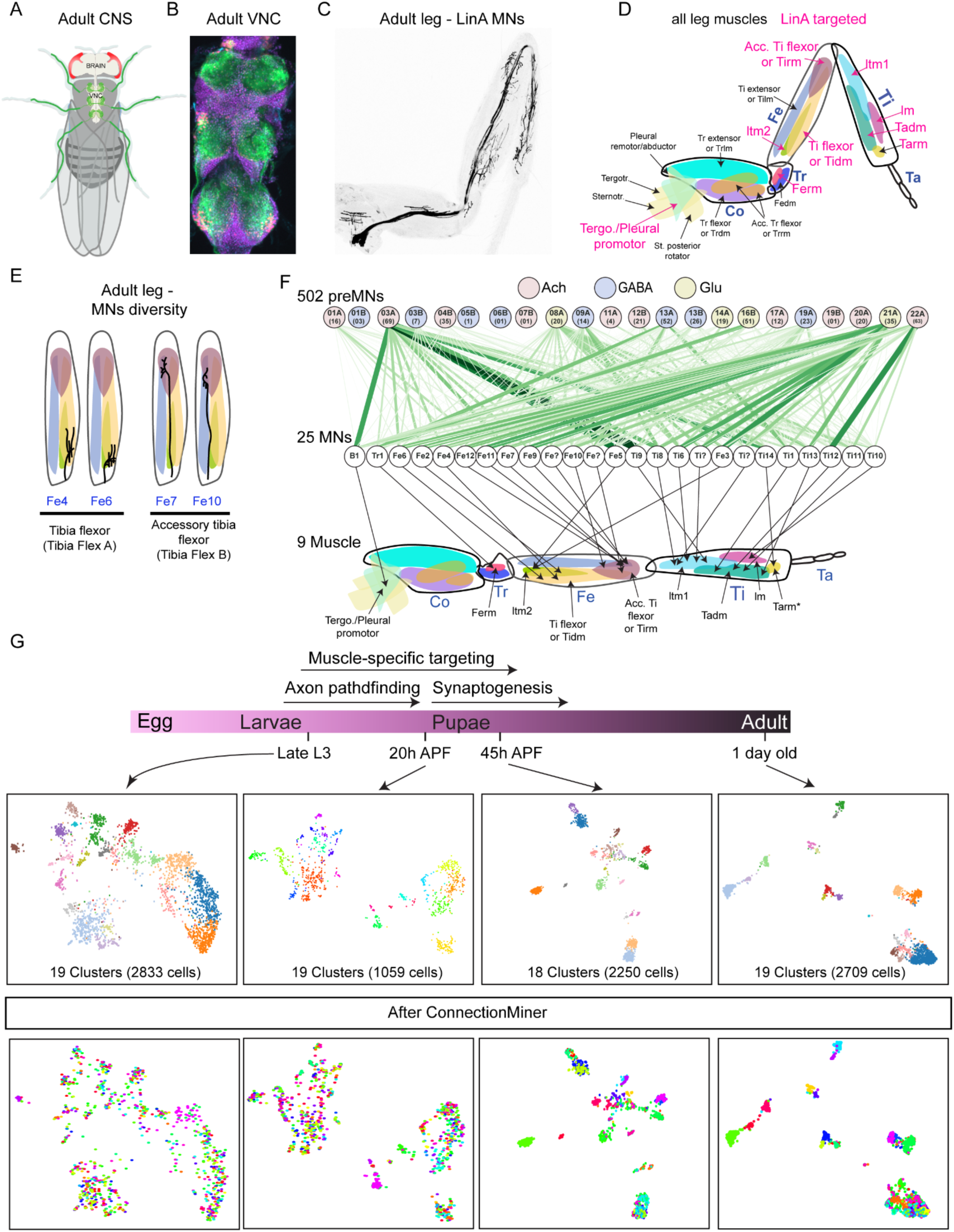
Transcriptional atlas of LinA-born MNs and their connectivity with preMNs. (A) Location of Drosophila leg MNs in the CNS. (B) Confocal image of LinA-derived genetically labeled both MN cell bodies and glia. (C) T1 leg showing the 29 MNs generated by LinA. (D) Schematic of all leg muscles in 5 leg segments (coxa, trochanter, femur, tibia and tarsus). Leg muscles in pink are those targeted by the 29 LinA MNs. (E) Schematic of two LinA-derived MNs that target the Tibia flexor and Accessory Tibia flexor muscles, with their axon morphologies schematized. (F) *Top row*: the neural stem cell hemilineages that give rise to the 701 local preMNs that synapse onto the 29 LinA MNs (*middle row*); *bottom row*: the leg muscles LinA MNs target. Numbers in the hemilineages circles indicate the number of LinA-contacting preMNs generated by that lineage and the color depicts the predicted neurotransmitter. The thickness of green lines depicts the proportion of total preMN inputs on a MN as seen in FANC. (G) Approximate time frames of the different steps in the leg MN development (top) and UMAP visualizations of scRNAseq dataset at four developmental stages before (*middle*) and after using ConnectionMiner (*bottom*).

## Results

### Transcriptomes of postembryonic LinA-born leg MNs

The adult leg MNs of Drosophila have their cell bodies and dendrites in the adult ventral nerve cord (VNC; Figure 1A, B) and axons that target specific muscles in each adult leg (Figure 1C, D). Of the ∼69 total leg-targeting MNs, 29 are derived from LinA and these target 9 of 18 leg muscles (Figure 1C, D). In addition to their stereotyped muscle targeting, each LinA MN is born in a stereotyped birth order and has a stereotyped dendritic arborization pattern (Baek & Mann, 2009; Guan et al., 2022). Despite the unique birth order of LinA MNs, several target the same muscle with similar arborization patterns (e.g., Figure 1E), underscoring the challenge of solving the neuronal wiring problem.

In addition to having a complete description of MN morphologies, recent electron microscopy (EM) reconstructions of the VNC have identified most of the synaptic connections by preMNs, which directly synapse onto MNs (Lesser et al., 2024; Cheong et al., 2024). These preMNs, mainly local interneurons, provide about 60% of the total input into MNs and are generated from 22 different hemilineages ((Lesser et al., 2024); Figure 1F). The goal of this study is two-fold: first, to document the molecular diversity of individual MNs as they mature, and second, to reveal the molecular components that specify the stereotyped connections between these three layers of the leg MN circuit – preMNs–MNs–leg muscles.

To identify the genes expressed in individual MNs, we carried out single cell RNA sequencing (scRNAseq) of the 29 MNs generated by LinA/15B neuroblasts at four developmental time points using the 10X Genomics platform (Figure 1G). We used a fluorescent reporter immortalization tool (Awasaki et al., 2014) to label all MNs born from the LinA neuroblast and FACS-sorted MNs at four developmental time points: 1) late third instar larvae (L3) when the MNs are postmitotic and axons have targeted the leg imaginal discs; 2) 20 hrs after puparium formation (APF), when 2° axon branches elaborate, 3) 45 hrs APF, when 3° branching initiates; and 4) one-day old adults, after MN maturation has completed ((Venkatasubramanian et al., 2019))(Figure 1G). We sequenced 7208 cells at L3, 3099 cells at 20 hrs APF, 3853 cells at 45 hrs APF and 4566 cells at the adult stage, with median unique molecular identifiers (UMIs) of 11,714, 16,020, 14,938 and 6,288, respectively.

After quality control and filtering for MNs based on VGlut expression, we performed unsupervised clustering on our scRNAseq data using the Phenograph implementation of the Louvain community detection method (Levitin et al., 2019). This resulted in 2833 cells for late L3, 1059 cells for 20 hrs APF, 2250 cells for 45 hrs APF and 2709 cells for the adult, which form 19, 19, 18 and 19 clusters, respectively (Fig. 1G). Visualizations of these cells and clusters were generated by two-dimensional uniform manifold approximation and projection (UMAP) plots (Figure 1G). We confirmed that all sequenced cells are derived from the LinA neuroblast based on the expression of genetic markers (nls-tdTomato-sv40 and myr-GFP-sv40; Figure S2) and known MN markers, including the presence of VGlut and the absence of Repo (a glia marker; Figure S2). In total, these scRNAseq datasets comprised 8851 cells that represent approximately 300x coverage of each LinA-born MN.

### Using EM connectomes to resolve scRNAseq cluster assignments

For the initial annotation of late L3 clusters, we used a reference dataset of morphological TFs (mTFs) known to be expressed in immature MNs ((Guan et al., 2022); Figure S3A). Additionally, we performed immunostaining against proteins that are predicted to be sparsely expressed in transcriptional clusters along with known mTFs. For example, as shown in Figure S3B, we found co-expression of Nubbin (Nub) and Kruppel (Kr) in one MN, indicating that transcriptional cluster 16 labels Tr1, an early-born MN. Similarly, we found co-expression of the late-born MN marker Prospero (Pros) with POU domain motif 3 (Pdm3), suggesting that late L3 clusters 0, 1, 2, 3, 7, 9, 15 and 17 correspond to late-born MN clusters (Figure S3C). This approach allowed us to identify 8 of the 19 transcriptional clusters at the late L3 stage (Figure S3D).

For the initial annotation of adult scRNAseq clusters, we used T2A-split-Gal4 (Chen et al., 2023; Lee et al., 2018) or MiMIC-based T2A-Gal4 lines to characterize the expression of genes or gene pairs predicted to be sparsely expressed in these clusters. For example, as shown in Figure S3E, using T2A-split-Gal4 for gene Tj or Salm along with VGlut, we found that adult cluster 12 labels Ti11 MN based on its muscle targeting and axon branching morphology (Figure S1). Using this strategy, we annotated 14 out of 19 clusters at the adult stage, which allowed the assignment of 16 of 29 LinA-derived MNs (Figure S2F).

The standard approach for analyzing scRNAseq data described above discovered fewer than the expected 29 MNs, suggesting that morphologically similar MNs may be difficult to distinguish using only transcriptome data. We reasoned that additional information, such as the connectome, could help annotate and resolve scRNAseq data. Building on this idea, we developed a machine learning algorithm, ConnectionMiner, which leverages the connectivity matrix between preMNs and MNs and preMNs and preMNs and integrates this information with transcriptome data of our partially annotated MNs and adult VNC (Figure 2) (see Methods for details). The idea is that MNs with similar morphologies may receive distinct preMN inputs, which can be used to distinguish them from a set of sequenced cells. For example, a T2A-split-Gal4 reagent that labels adult clusters 2 and 3 (Foxo ∩ Tey) is expressed in four MNs in the adult (Figure 3A), suggesting that these clusters are not fully resolved into individual MNs. Moreover, Fe10 and Fe11 target the same muscle in the femur and have very similar transcriptomes, but their inputs from preMNs differ significantly (Figure 3B). In addition to leveraging differences in their connections, ConnectionMiner can incorporate additional constraints, such as information about muscle targeting, the initial annotation of adult clusters, and MN birth order, which is well-defined for all LinA MNs. Using this tool, cells present in the annotated as well as unannotated clusters were reclustered to assign cells to a MN (Figure S4). After the application of ConnectionMiner, adult cluster 2 is correctly split into two, one for Fe10 and one for Fe11, and cluster 0 resolves into 10 MNs (Figure 3D). We also applied ConnectionMiner to the published scRNAseq dataset for the adult VNC that contains neuroblast (NB) hemilineage information from which all preMNs are born (Allen et al., 2020; Soffers et al., 2025). Notably, ConnectionMiner was also able to resolve these transcriptional clusters into individual preMNs (Figure 2).

**Figure 2:**
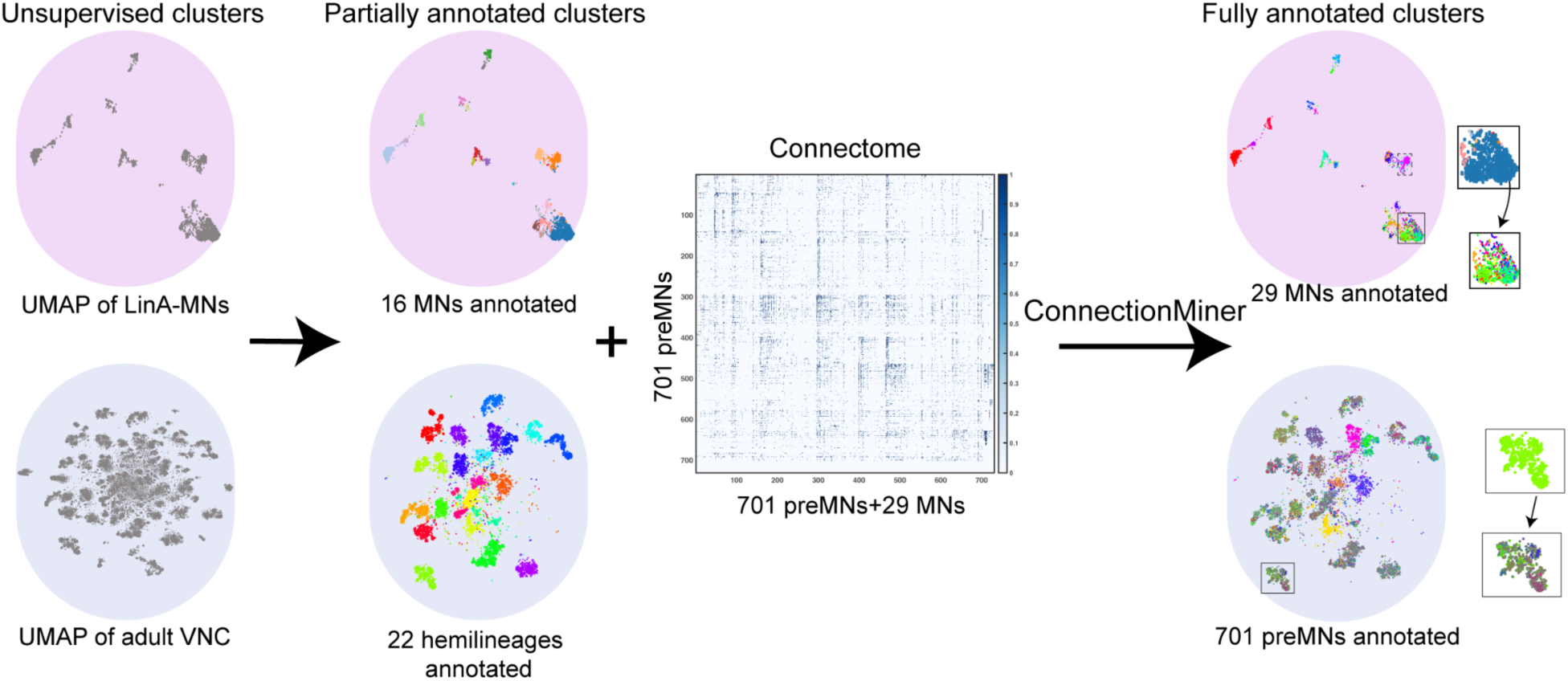
Overview of ConnectionMiner. The left pair of UMAPs are the result of unsupervised clustering for the 29 LinA MNs (top) and their connected preMNs (bottom). These clusters were partially annotated using lineage information and genetic reagents (middle UMAPs). ConnectionMiner further resolves these partially annotated clusters with the by inputting the connectivity matrix of preMNs and MNs to generate a fully refined set of clusters (right pair of UMAPs). The insets highlight ‘before’ and ‘after’ annotations for both MNs (top) and preMNs (bottom).

**Figure 3:**
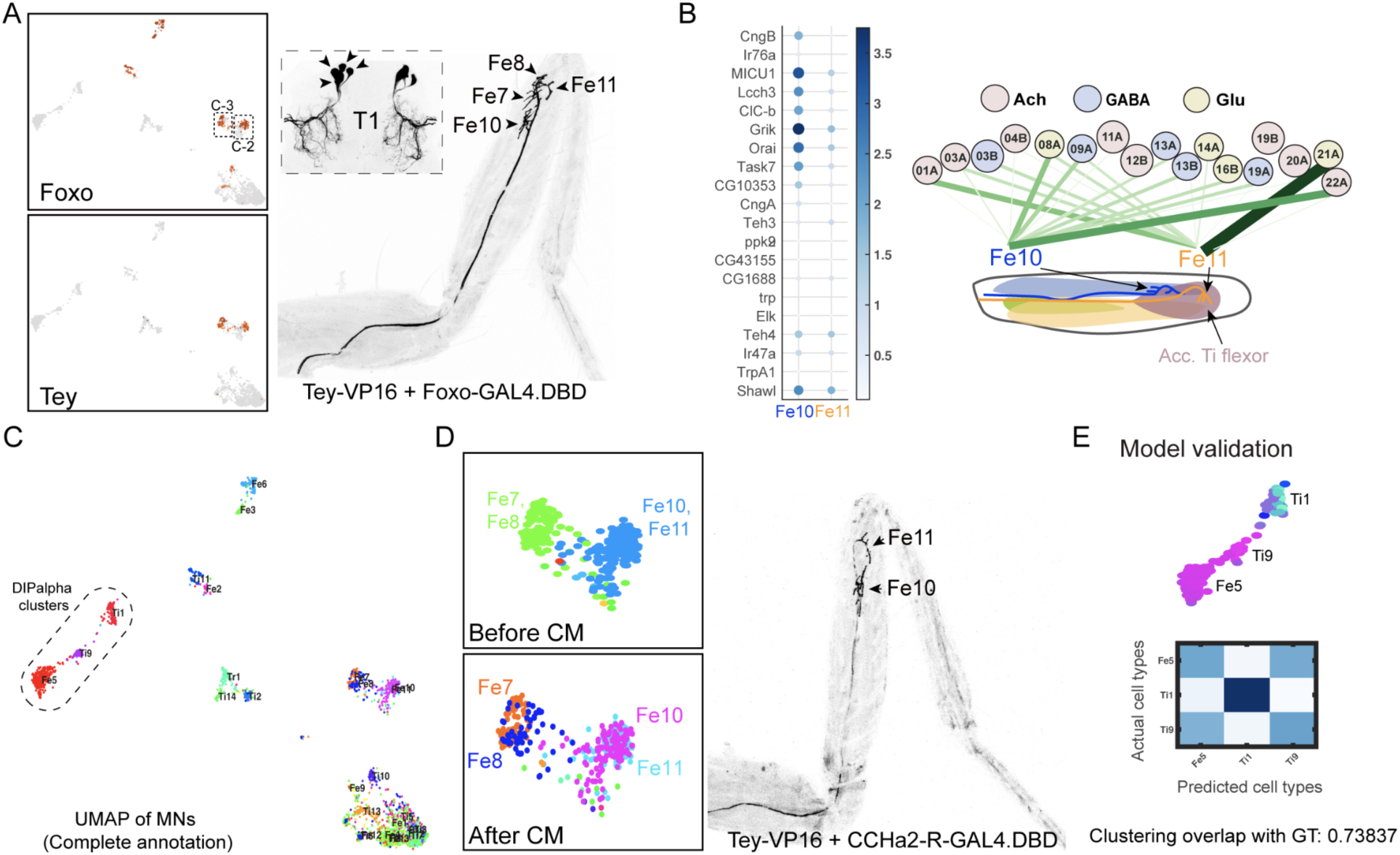
Tests of ConnectionMiner. (A) Split-GAL4 for genes *foxo* and *tey,* which label only two clusters, is expressed in 4 leg MNs as evident from 4 MN cell bodies detected in each VNC hemisegment. (B) Heatmap showing very similar differentially expressed genes for two MNs, Fe10 and Fe11 (left), yet these MNs are connected by different sets of preMN (right). (C) UMAP visualization of adult scRNAseq cluster after ConnectionMiner resolved into 29 clusters. The three DIP-alpha clusters are highlighted. (D) ConnectionMiner correctly split clusters 2 and 3 into four (left) and predicted split-GAL4 of CCHa2-R and tey would be expressed in two MNs (Fe10 and Fe11; right). (E) We further tested ConnectionMiner by determining if it could correctly reassign the three DIP-alpha MNs after removing their assignments. The heat map compares the new cell assignments (horizontal axis) with the ‘ground truth’ (vertical axis), revealing an accuracy of 0.74.

Once the ConnectionMiner model is generated, we ask whether the refined assignments of cells to neurons are correct. Indeed, this is the case. For example, ConnectionMiner correctly determined that several clusters in our MN scRNAseq dataset corresponded to multiple MNs (Figure 3A,D). Further, ConnectionMiner correctly split those clusters into individual MNs and correctly predicted new markers for labeling two of these four MNs (Figure 3D).

A second test of ConnectionMiner was to remove all prior annotations and cell assignments for a subset of MNs and determine if the algorithm could correctly reassign these cells to the correct clusters. We tested this for the three leg MNs that express the Ig domain CAMs, DIP-ɑ. After removing all prior information for these cells, the algorithm was able to reassign them to the correct MN with an accuracy of 74% (Figure 3E). Thus, although its performance is imperfect, ConnectionMiner is a valuable tool that can help refine ambiguous transcriptome clusters into higher resolution clusters. Further, because the algorithm accomplishes this using gene expression data, one output are sets of molecules that are predicted to drive these refined cluster annotations.

### preMN connectivity correlates with transcriptome diversity of MNs

Among the ∼69 leg targeting MNs in each thoracic hemisegment, including the 29 derived from LinA, most target distinct sets of muscle fibers and have different axon and dendritic morphologies, patterns of connectivity, and physiological properties (Baek & Mann, 2009; Azevedo et al., 2020; Lesser et al., 2024; Figures 1D, 1F and S1). On the other hand, some leg MNs have very similar morphologies, consistent with their similar transcriptomes (e.g., Figure 3B). To determine to what extent these features are reflected in their transcriptomes during development, we generated force-directed lineage trajectories of the 29 LinA MNs from the late L3 to the adult and then used the optimal transport method (Schiebinger et al., 2019) to link clusters between the four time points. The resulting trajectories allow us to follow the development of all 29 LinA MNs from late L3 to the adult (Figure 4A and S5A).

**Figure 4:**
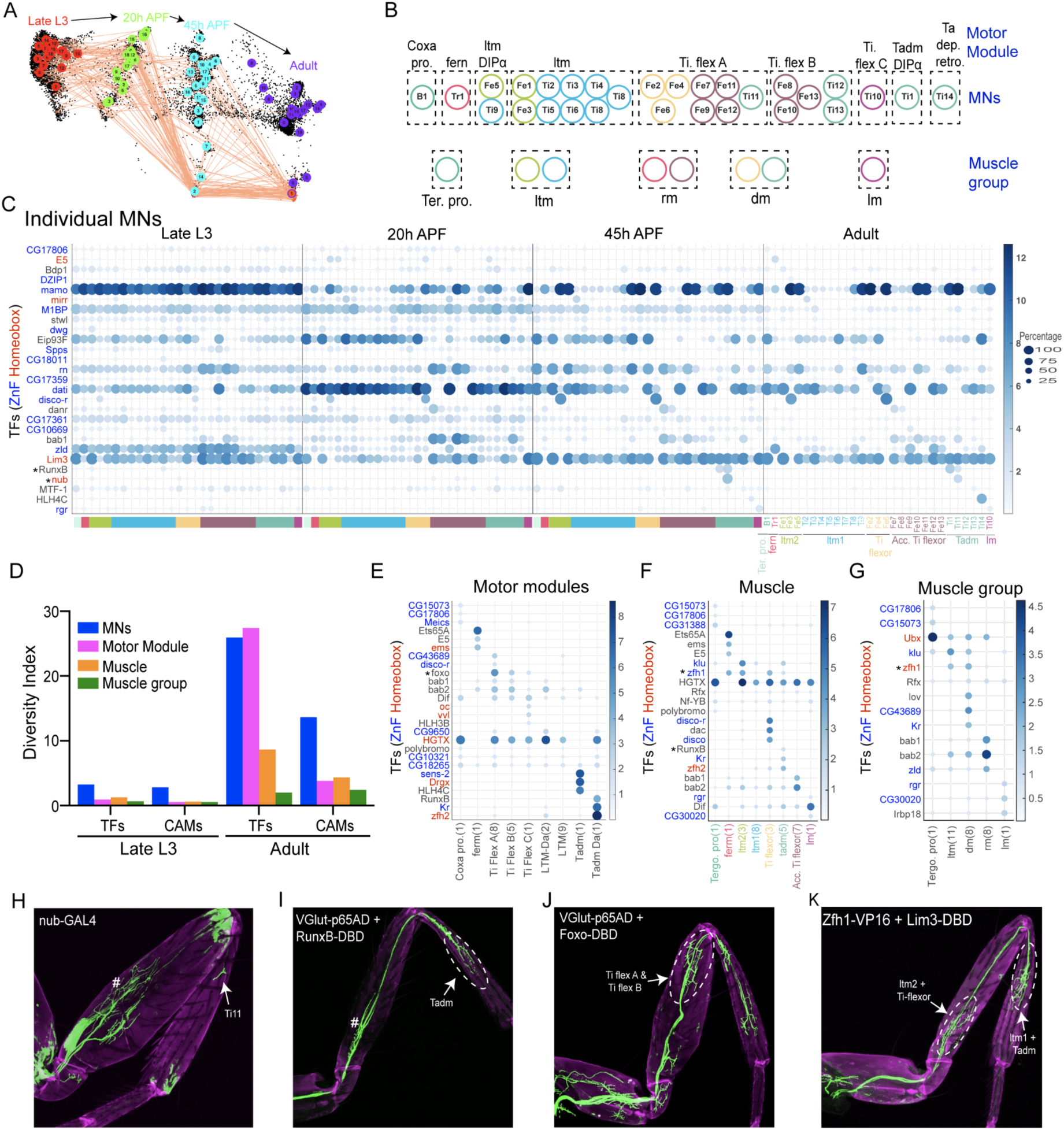
Motor modules and TFs correlate best with MN diversity signatures. (A) Optimal transport method showing linkage between clusters across four time points. Fe5-ltm MN is highlighted as an example. (B) Grouping of MNs into different categories - motor modules and muscle group are shown. Colored circles around MNs show their muscle target, as depicted in panel C. (C) Heatmaps showing top 1 TF expressed in 29 leg MN in adults and their developmental expression pattern in other time points.Note that the patterns become more specific as MNs mature. ((D) Diversity index showing the fraction of genes belonging to TFs and CAMs gene families at Late L3 and adult stage, calculated for individual MN as well as for different MNs grouping. (E-G) Heatmaps showing expression of top 3 TFs in MNs when grouped according to motor modules (E), muscles (F) and muscle group (G). (H-K) T1 leg showing expression of few selected genes from panel C-G. * indicates validated gene.

To characterize the transcriptomes in each of the 29 LinA-derived MNs, we used a set of 2595 highly variable genes (HVGs) that were determined to be differentially expressed among these MNs (Figure S5B). These HVGs represent 34% of the total expressed genes in these MNs. The two most variable gene families are the TFs and cell adhesion molecules (CAMs), which together comprise 15.5% of the HVGs (Figure S5B). To quantify the degree to which these and other gene families are differentially expressed over time, we calculated a parameter called the Diversity Index (DI). The DI is calculated by carrying out a multivariate analysis of variance (MANOVA) to determine the amount of variation that exists between each MN for any group of genes (see Methods). Using this measurement, we found that TFs and CAMs are the most differentially expressed gene families at all time points, with peak DI values at 45h APF and the adult (Figures 4D and S5C). When subfamilies of TFs and CAMs were examined, we found that zinc-finger (ZnF) and homeodomain (HD) TFs had the greatest diversity (Figure S5D), while for CAMs immunoglobulin (Ig) domain proteins had the greatest diversity (Figure S5E). A similar picture emerges when we examine which genes differ the most between individual MNs (Figures 4C and S6): variability, particularly for ZnF, HD, and Ig domain genes, is most obvious at 45h APF and the adult. These trends remain the same even when we search for top differentially expressed genes (DEGs) at each time point (Figure S7). However, the specific genes identified differ depending on the time point being analyzed, suggesting that different combinations of TFs and CAMs are being used at different times during development. We validated a subset of these predictions (indicated by ‘*’ in Fig. 4C) for genes expressed in a subset of MNs (e.g., *nub* and *RunxB* (Figure 4H-I)).

Leg MNs can be grouped in a number of ways (Figure 4B). For example, one grouping depends on which muscles each MN targets. The 29 LinA MNs target 8 different leg muscles. These 8 leg muscles can be reduced to 5 different **muscle groups**, such as those targeting a ‘depressor’ or a ‘reductor’ muscle. Third, the 29 LinA MNs can be grouped according to the preMN inputs they receive, referred to as **motor modules** (Lesser et al., 2024). Accordingly, the LinA MNs fall into 9 distinct motor modules (see Methods for a complete list of each group). In principle, these different groupings might be reflected in their transcriptomes; i.e., MNs that target the same muscle or are within the same muscle group might have more similar transcriptomes than MNs in different groups. When we calculated the Diversity Index for each of these groups, we found that motor modules had the highest values, particularly for transcription factor gene families (Figures 4D-G, S8). We validated a subset of these predictions (indicated by ‘*’ in Figure 4E-G) for genes expressed in a subset of motor modules, muscle or muscle groups e.g., *foxo* (for Tibia Flex A and Tibia Flex B) and *zfh1* (for long tendon muscle (ltm) 2, ltm1, Tibia flexor, and Tarsal depressor muscle (tadm); Figure 4J-K)).

Notably, we found that TFs were the only gene family that stood out for each MN and motor module group. These observations suggest that TFs are the main drivers of MN diversity. Further, they suggest that MNs within the same motor module tend to share similar transcriptomes compared to MNs in different modules. Further, similarities between transcriptomes are less pronounced when MNs are grouped according to their other properties, such as which muscle they target. These observations suggest that connectivity to preMNs correlates best and may be a driving force for generating MN transcriptional diversity.

### Computational inference with ConnectionMiner identifies candidate synaptic protein interactions between preMNs and MNs

Once ConnectionMiner is used to derive a full annotation of all 701 preMNs and 29 MNs, we can use it to calculate average gene expression profiles for each cell type (see Figure S4D). Then, using this information and the known connectome, we sought to identify specific gene-pair combinations most predictive of synaptic connectivity between preMNs and MNs (Figure 5). Here, we tested this for the preMNs that interact with 29 LinA-derived leg MNs and the refined scRNAseq dataset for the adult fly VNC, along with preMN-MN and preMN-preMN connectivity datasets (Figure 2). First, for each pair of neurons in the connectome, we computed a co-expression feature for every possible pair of genes in the transcriptome. By comparing gene co-expression levels in connected versus non-connected neuron pairs using a univariate rank-sum test, we measured an effect size (how strongly each gene pair distinguishes connected from non-connected pairs) and a statistical significance (p-value). This is analogous to the method presented in Taylor et al.(Taylor et al., 2021). This is visualized in Figure 5A, where we show a “volcano” plot of effect size (horizontal axis) versus statistical significance (vertical axis) for each gene pair. Gene pairs lying toward the right and with higher −log10(p) values (i.e., points in the upper-right region) are most strongly associated with connectivity. Notably, we observed that a relatively small fraction of gene pairs showed both high effect sizes and strong significance, suggesting that the connectome may rely on a limited combinatorial code of genes to drive neuron-neuron synaptic specificity.

**Figure 5:**
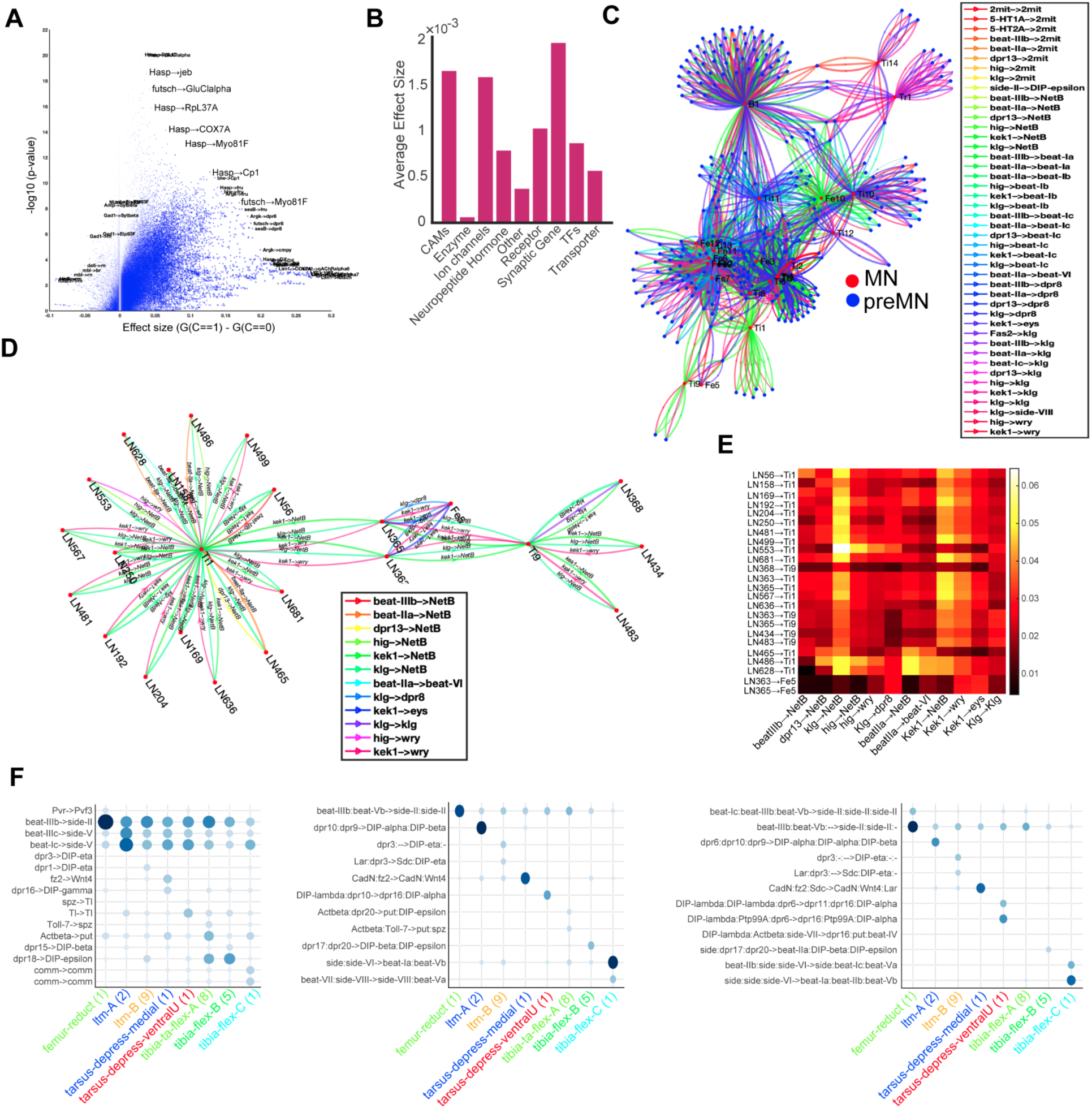
Identifying and visualizing gene-pair contributions to neuronal connectivity. (**A**) A volcano plot compares gene-pair effect sizes (horizontal axis) with their statistical significance (−log10 p-value, vertical axis) between connected and non-connected neuron pairs. Each point represents a gene-pair co-expression feature; points to the right with high −log10 p-values are candidate drivers of connectivity. (**B**) A bar chart shows the average effect size for significant gene-pairs grouped by their annotated gene family (e.g., transcription factors, cell-adhesion molecules), highlighting families most associated with strong connectivity effects. (**C**) A network diagram depicts the preMNs (blue nodes) and MNs (red nodes), with edges colored according to the top three gene pairs (ranked by effect size and probability) implicated in each connection. Each color corresponds to a distinct gene pair, illustrating the combinatorial molecular code underlying individual neuronal connections. The width of edges indicates the effect size of the gene pair multiplied by their co-expression probability in their corresponding neuron pair. (**D**) Zoom panel of a subnetwork of the preMN to MN connectome, showing the input connections for MNs, Ti1, Fe5 and Ti9 is shown. Edges are further labeled with text to indicate the gene pairs. (**E**) A heat map displays the product of effect size and co-expression probability for each gene pair (columns) in different presynaptic–postsynaptic neuron pairs (rows). Warmer colors (yellow/red) indicate gene pairs that are strongly predictive of synaptic connectivity for those specific neuron pairs, whereas cooler colors (black) represent weaker or no association. (**F**) Rows indicate differentially expressed gene pairs in different motor modules in the columns. The three plots show gene combinations that consider 1×1 gene pairs (left), as well as 2×2 (middle) and 3×3 (right), indicating the increasing levels of differential coding of motor modules with higher order gene combinations.

Since this initial analysis was done in an unbiased manner across all genes, we sought to group the effect sizes of gene pairs by their gene families. Specifically, identification of interacting molecules involved in neuronal wiring between preMNs and MNs, we generated a curated list of cell adhesion molecules (CAMs) and synaptic genes that are known from the literature to interact with each other (see Methods for details;(Battistini & Tamagnone, 2016; Carrillo et al., 2015; Li et al., 2017; Özkan et al., 2013; Sanes & Zipursky, 2020; Sergeeva et al., 2020; Sullivan & Bashaw, 2023; Yazdani & Terman, 2006)). Furthermore, we also classified genes into enzymes, ion channels, transporters, and neuropeptides as additional controls. Grouping these significant gene pairs by their gene families revealed that TFs, CAM, and synaptic-related genes contributed disproportionately to high effect sizes (Figure 5B). This result aligns with our previous observations that TFs and CAMs are the two gene families most diversified across MNs and preMNs. It also further underscores the notion that multiple molecular classes act in concert to define neuronal wiring.

To illustrate how specific gene pairs might combine to mediate connectivity, we generated a force-directed network diagram of all preMN (blue nodes) to MN (red nodes) edges (Figure 5C). Each directed edge from a preMN to a MN is replicated up to three times (colored lines), corresponding to the top three gene pairs implicated in that connection. Each gene pair is assigned a distinct color, and the edge thickness is proportional to the product of the pair’s effect size and its co-expression level in the corresponding neurons. This representation highlights how different neuron pairs may rely on partially overlapping sets of gene pairs to establish synapses. A zoomed-in subnetwork (Figure 5D) shows, for example, how the MNs defined by DIP-ɑ expression, Ti1, Fe5, and Ti9, each receive inputs from overlapping but not identical sets of preMNs, often distinguished by unique gene-pair combinations.

Within this set of DIP-ɑ expressing MNs, Ti1, Fe5, and Ti9, we further explored the combinatorial usage of gene pairs with a heat map (Figure 5E) in which each row represents a distinct preMN→DIP-ɑ expressing MN connection, and each column corresponds to one of the top implicated gene pairs. Warmer colors (yellow and red) indicate high ConnectionMiner scores (effect size × co-expression probability), whereas cooler colors (black) indicate weaker associations. The diagonal “blocks” of bright signals suggest that particular gene pairs strongly characterize specific connections, consistent with a modular pattern of usage. This heatmap also serves as a platform to design validation experiments since the score in each entry reflects the probability of observing a particular effect size by the indicated gene pair in the particular synapse, thereby enabling prioritization of knockout experiments.

Finally, we evaluated whether higher-order gene combinations (e.g., pairs of pairs, or triplets of genes) are more predictive of differential connectivity patterns. Because we found that motor modules, which are based on preMN inputs, correlate best with distinct MN transcriptomes (Figure 4E), we simplified the problem by grouping the LinA-connected preMNs into 9 premotor modules: coxa promote, femur reductor, tibia flex A, tibia flex B, tibia flex C, ltm DIP-ɑ, ltm, tarsus retro depressor and tarsus depressor DIP-ɑ (Lesser et al., 2024). We then used ConnectionMiner annotations to find distinct gene co-expression patterns observed in the connections in specific preMN modules that connect with MN modules (Figure 5F, S9). Importantly, we found that the combinatorial expression of two or three CAMs (as opposed to single CAMs) provided the most distinct predictions, as evidenced by the observed diagonal pattern of differential co-expression of the gene combinations. Specifically, in Figure 5F, we compared how 1×1 gene pairs (left), 2×2 gene combinations (middle), or 3×3 gene sets (right) distinguish the connections observed across motor modules. Indeed, adding additional genes to the combinatorial code (i.e., 2×2 or 3×3 sets) yielded further differentiation among modules, highlighting that multiple interacting gene sets are likely required to achieve robust specificity in this circuit. These findings argue that there is a combinatorial code of CAMs that is required for preMN-MN connectivity.

Together, these analyses illustrate how ConnectionMiner pinpoints both individual genes and gene-pair combinations implicated in preMN-to-MN connectivity. The results strongly support a model in which a limited set of cell-adhesion and transcriptional regulators applied in diverse combinations in heterophilic/homophilic manners mediate the precise assembly of the adult leg motor circuit.

## Discussion

In this study, we provide an in-depth analysis of the molecular steps that allow a set of immature post-mitotic MNs to mature into distinct MNs, with stereotyped axon and dendritic morphologies and distinct connections to upstream preMNs and to downstream muscles. To facilitate this analysis, we developed a novel machine learning tool, ConnectionMiner, that can be used to decode neuronal wiring. We applied this tool to dissect three layers of the adult leg motor circuit – preMNs to MNs to muscles. ConnectionMiner integrates connectomics and transcriptomics datasets along with leveraging other parameters like birth order and muscle targeting to resolve ambiguous identities of neurons with similar morphologies and transcriptomes. Additionally, ConnectionMiner predicts the molecular code that drives connectivity. Importantly, our computational tools bridge the gap between single-cell transcriptomics and connectomics, thereby helping to decode neuronal wiring.

### ConnectionMiner resolves transcriptional ambiguity between similar MNs

It is widely accepted that scRNAseq can delineate neuronal diversity based on transcriptional heterogeneity observed due to morphological differences in neurons (Kurmangaliyev et al., 2020; Özel et al., 2021). However, using conventional approaches that examine the expression patterns of cluster markers or sparsely expressed genes, we were only able to partially annotate adult scRNAseq clusters. Interestingly, we found that LinA-derived MNs, which share similar morphology and innervate the same leg muscles for example, those targeting the accessory tibia flexor or long tendon muscles— were poorly annotated. This observation suggests that these MNs may have highly similar transcriptome profiles that are difficult to discriminate using only transcriptomic data. Previous studies have identified transcriptional heterogeneity in neuronal subtypes with similar morphology but distinct synaptic specificities, such as Dm8 neurons (Courgeon & Desplan, 2019; Menon et al., 2019), indicating that neuronal connectivity can serve as a key factor in resolving transcriptional heterogeneity. Surprisingly, in our study, we found that a group of MNs with highly similar transcriptomic profiles also received similar preMN inputs and innervated the same leg muscles, making them transcriptionally indistinguishable using conventional methods. We anticipated these challenges, as these MNs share a similar combinatorial morphological transcription factor (mTF) code (Guan et al., 2022).

To address this issue and identify transcriptional heterogeneity within these MNs, ConnectionMiner integrates transcriptomic data from both MNs and preMNs, along with connectivity information from preMNs-MNs and preMNs-preMNs derived from EM-connectomics datasets. ConnectionMiner successfully resolved and annotated adult MN scRNAseq clusters for groups of MNs with highly similar transcriptomic profiles. We validated this new adult scRNAseq cluster annotation using T2A-split-Gal4 lines for genes predicted to be expressed in these MNs. In addition to analyzing MN scRNAseq datasets, ConnectionMiner was also applied to published scRNAseq datasets of the adult VNC, which contain transcriptomic information for each hemilineage from which these preMNs originate (Allen et al., 2020; Soffers et al., 2025). Each hemilineage has distinct morphological and neurotransmitter identities (Lacin et al., 2019), which were used in EM connectomic datasets to classify and group preMNs into their respective hemilineages. Notably, MNs receive synaptic input from preMNs from multiple hemilineages (Lesser et al., 2024). However, preMNs exhibit stereotyped synaptic connectivity with groups of MNs involved in synergistic motor functions, such as flexion. Leveraging this information, we applied ConnectionMiner and successfully identified individual preMNs that provide synaptic input to the 29 LinA MNs at the transcriptional level.

These findings demonstrate that if connectivity data are available, ConnectionMiner can be deployed to decode transcriptional heterogeneity among morphologically and functionally similar neurons. Furthermore, ConnectionMiner is capable of resolving transcriptional differences in both newly generated and publicly available scRNAseq datasets, overcoming the limitations of traditional clustering algorithms in distinguishing neuronal subtypes with highly similar transcriptomic profiles.

### TFs drive three layers of the leg motor circuit formation

The formation of the three layers of the leg motor circuit, i.e., preMN - MN - leg muscle, is a stepwise process during development. It involves neuronal specification, axon guidance and synapse formation, neuromuscular junction formation and preMN-MN circuit assembly. Previous studies on LinA MNs have shown that combinations of TFs expressed early in their maturation are critical for their morphological identity (Guan et al., 2022). Our analysis emphasizes the importance of TFs in driving MN diversity. Irrespective of how MNs are grouped into different categories, we found that the Diversity Index is consistently higher for TFs compared to other functional gene families. Moreover, we found that when grouped into motor modules, MNs have the highest Diversity Index value compared to groupings based on muscle or muscle type. These observations suggest that preMN inputs correlate best with MN diversity. Further, among the various families of TFs, we found that Zn-finger and homeobox TFs appear to play the most important role in driving diversity.

### ConnectionMiner identifies candidate synaptic protein interactions between preMNs and MNs

For the leg motor circuit to function properly, leg MN dendrites must receive synaptic input from the correct subsets of preMNs present within the leg neuropil (Lesser et al., 2024). Importantly, for the precision of preMN connectivity, each preMN must identify the target motor modules during development and provide proportional input that depends on MN size. This ensures that all MNs within a module are coordinately excited or inhibited in response to changes in the activity of each preMN. Similarly, sets of preMNs synapse onto sets of MNs that execute similar functions, for example, flexion (Lesser et al., 2024). This pattern of neuronal wiring between preMNs and MNs follows Hebb’s rule: “fire together, wire together” (Stent, 1973; Hebb, 2005). To achieve this robustness, molecular interactions between two interacting neurons or cells are likely to play a crucial role in establishing their connectivity. There are multiple examples where CAMs have been shown to play a role in connectivity. For example, pairs of interacting proteins of the Beaten path (Beats) and Sidestep (Sides) families play a role in synaptic targeting in the visual system (Carrier et al., 2024; Yoo et al., 2023), while members of the DIP and Dpr Ig domain families have been shown to play a role in a subset of MNs to target the correct leg muscles (Venkatasubramanian et al., 2019; Morano et al., 2024; Lopez et al., 2024). However, there are also examples where removing a CAM results in no apparent phenotype (e.g., (Venkatasubramanian et al., 2019; Morano et al., 2024; Lopez et al., 2024). Negative results such as these suggest that, for many neurons, combinations of CAMs may be required. Consistent with this idea, the results from ConnectionMiner show that pairs and triplets of CAMs result in the greatest amount of discrimination between MNs (Figure 5F and S9).

### Comparisons to other cell-cell communication methods

Several computational methods have been developed to annotate cell types and infer cell-cell communication from single-cell transcriptomic data, yet each exhibits distinct limitations. For instance, CellChat (Jin et al., 2021) infers intercellular communication by leveraging curated databases of ligand-receptor interactions and permutation-based statistics; however, its predictions are constrained by the completeness and accuracy of these databases and it does not incorporate anatomical or connectivity data. Similarly, CellPhoneDB (Efremova et al., 2020) uses the expression of multi-subunit ligand-receptor complexes to predict potential interactions, but its framework is agnostic to the actual synaptic architecture and therefore may miss key functional connections. scConnect (Jakobsson et al., 2021) offers an exploratory approach to cell-cell communication analysis by integrating transcriptomic profiles, yet it, too, lacks the integration of precise connectomic data, which is essential for resolving functional synaptic specificity. In contrast, ConnectionMiner innovates by combining scRNAseq data with EM–derived connectomes within a unified factorization framework. This integration not only refines ambiguous transcriptomic clusters by incorporating known synaptic connectivity, but it also directly assigns probabilities to gene pair interactions that are predictive of synaptic strength. By leveraging both gene expression and structural connectivity information, ConnectionMiner provides a more robust and mechanistically informed model of neuronal wiring, thereby overcoming limitations inherent in methods that rely solely on transcriptomic or curated interaction data.

Traditional approaches for inferring the molecular rules governing neuronal connectivity such as those developed by Barabási et al. (Barabási & Barabási, 2020), Kovács et al. (Kovács et al., 2020) and Taylor et al. (Taylor et al., 2021) rely on fully confident, pre-annotated cell-type-specific gene expression profiles. These methods assume that the cell types and their corresponding transcriptional signatures are known a priori, and they use this static information to infer genetic connectivity rules. In contrast, ConnectionMiner is uniquely designed to operate on partially annotated datasets by jointly learning both probabilistic cell-type assignments and the underlying connectivity rules. In our framework, the connectivity rules are iteratively refined using the connectome data, and as these rules increasingly reflect the true wiring of the circuit, the cell-type annotations become more confident. This positive feedback loop enables ConnectionMiner to not only resolve ambiguous cell identities but also to robustly predict the combinatorial gene interactions that drive synaptic connectivity. Consequently, ConnectionMiner provides a more flexible and dynamic approach to deciphering the molecular code of neuronal wiring compared to previous static annotation–dependent methods.

### Comparisons to other cell type annotation methods

Cell type annotation methods such as SingleR (Aran et al., 2019) and CellAssign (Zhang et al., 2019) rely exclusively on transcriptomic signatures and well-curated reference datasets to assign cell identities. These approaches work well when high-confidence, distinct gene expression profiles for each cell type are available. However, in complex tissues or developmental contexts where cell types are only partially annotated or exhibit highly similar transcriptional profiles, these methods often fail to resolve ambiguities. In contrast, ConnectionMiner innovates by integrating connectomic data information about the actual synaptic connectivity between cells with single-cell gene expression. This additional layer of data allows ConnectionMiner to drive cell type annotation even when transcriptomic differences are subtle, creating a positive feedback loop: as connectivity rules are refined, cell type assignments become more confident, and vice versa. This integrative strategy enables the identification of connectivity-specific molecular codes that traditional, transcriptomics-only approaches cannot capture.

### Limitations of this study

Despite the promising results of ConnectionMiner, several limitations remain. The accuracy of our approach is intrinsically tied to the quality and completeness of the underlying connectome data. Although recent electron microscopy reconstructions provide unprecedented detail, weak or transient synapses may be underrepresented, potentially biasing the inferred connectivity rules. Also, scRNAseq data are subject to technical limitations such as dropout events and low capture efficiency, which can obscure the expression of low-abundance transcripts critical for precise cell type discrimination. Although ConnectionMiner does not require full annotations as traditional cell-cell communication methods do, it is greatly aided by partial annotations to constrain its solutions. The quality and completeness of the initial annotations have an effect on the downstream results that the method produces. Also, while we focus on curated gene family categorizations with special emphasis on CAMs, novel or uncharacterized genes involved in synaptic wiring may be overlooked. Similar to Kovacs et al. ConnectionMiner model’s assumption that synaptic connectivity can be linearly approximated by gene co-expression products, while computationally tractable, might oversimplify the complex, nonlinear biological interactions underlying synapse formation. Future versions of ConnectionMiner will incorporate non-linear decoder modules to overcome this potential limitation. Additionally, the probabilistic cell type assignment framework, although effective in resolving ambiguous clusters, has been primarily validated within the context of the Drosophila leg motor system, and its performance in other neural systems remains to be determined. Finally, while our computational predictions provide a valuable basis for hypothesis generation, extensive experimental validation is necessary to confirm the functional roles of the identified gene pairs and their combinatorial codes in establishing synaptic specificity.

### Conclusion

In this study, we introduce ConnectionMiner - a novel machine learning framework that integrates single-cell transcriptomics with connectomic data to resolve transcriptional heterogeneity through annotating distinct cell types and predicting candidate molecular interactions essential for neuronal wiring. By leveraging differential synaptic connectivity, ConnectionMiner resolves transcriptionally similar single cells into distinct neuron types and helps infer a combinatorial molecular code of connectivity. Although the predicted CAMs and interacting gene pairs that mediate connectivity await experimental validation, the iterative nature of our approach permits continual refinement as additional experimental data becomes available. Moreover, while our current implementation focuses on known interacting molecules, the framework is readily extendable to encompass all cell surface proteins, thereby offering the potential to discover novel molecular interactions governing circuit assembly. In conclusion, ConnectionMiner offers a new approach in decoding the molecular determinants of neural connectivity and provides a robust blueprint for future studies aimed at understanding the mechanisms of neural circuit formation across diverse systems and species.

## Materials and methods

### Key resources table

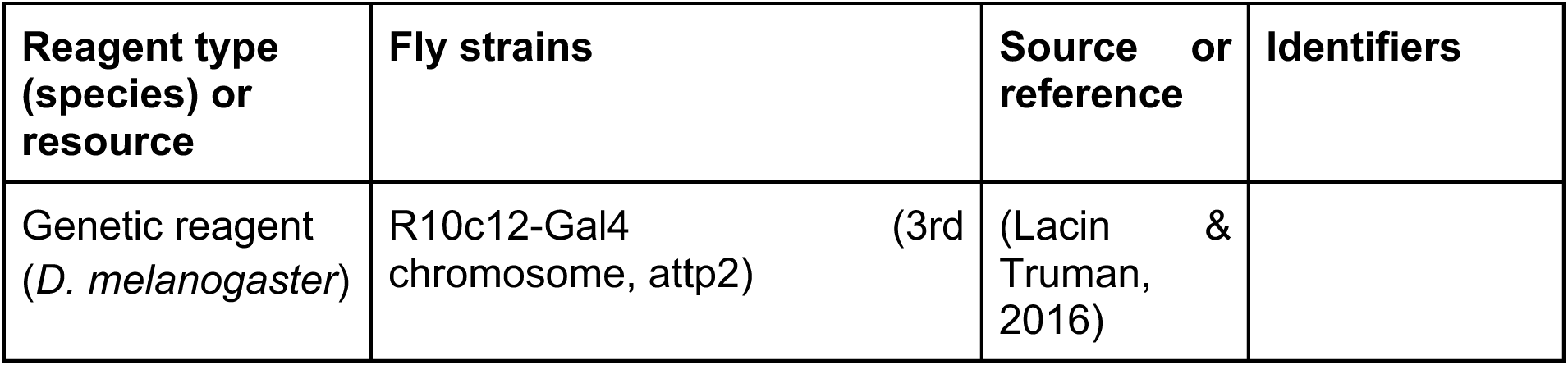
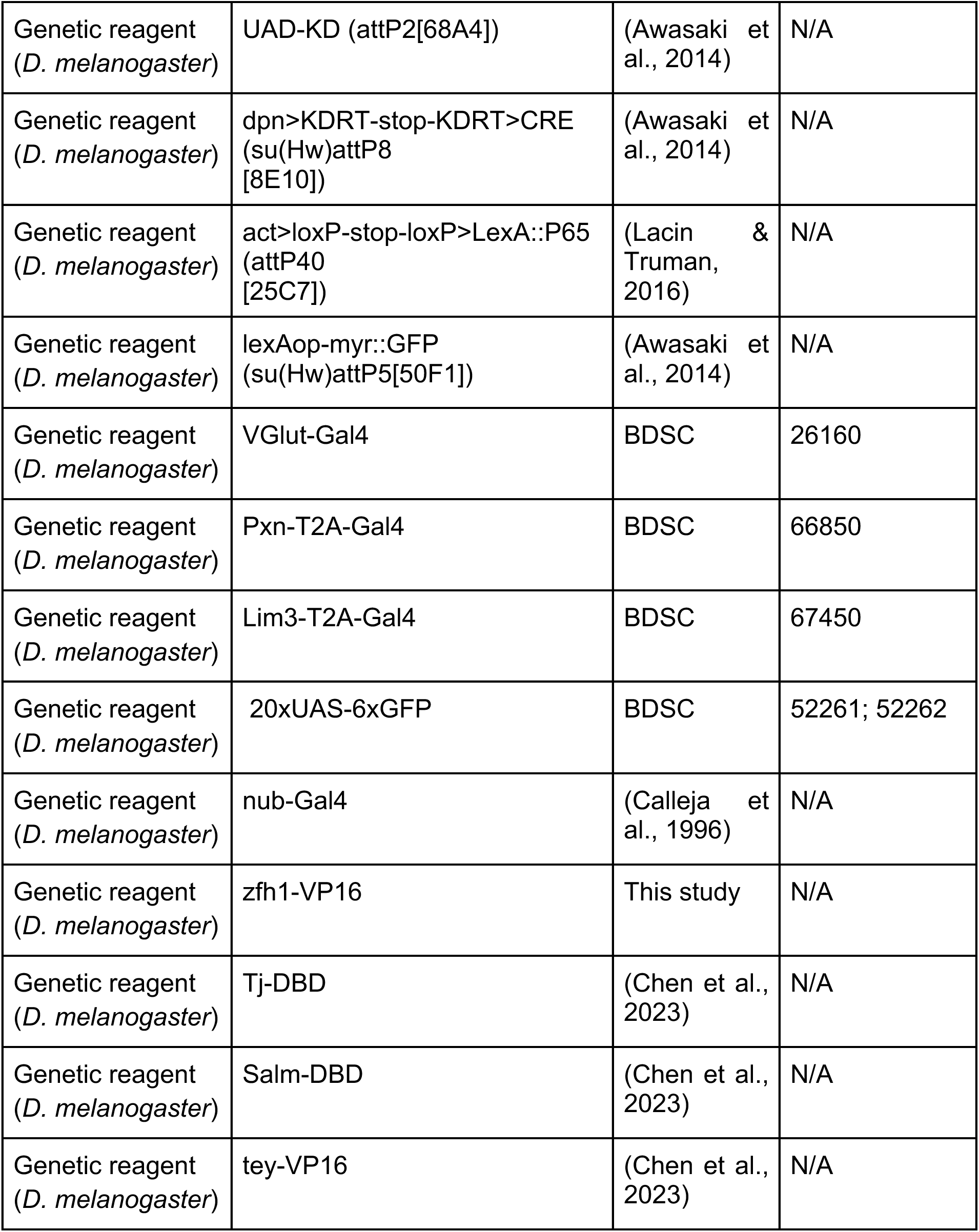
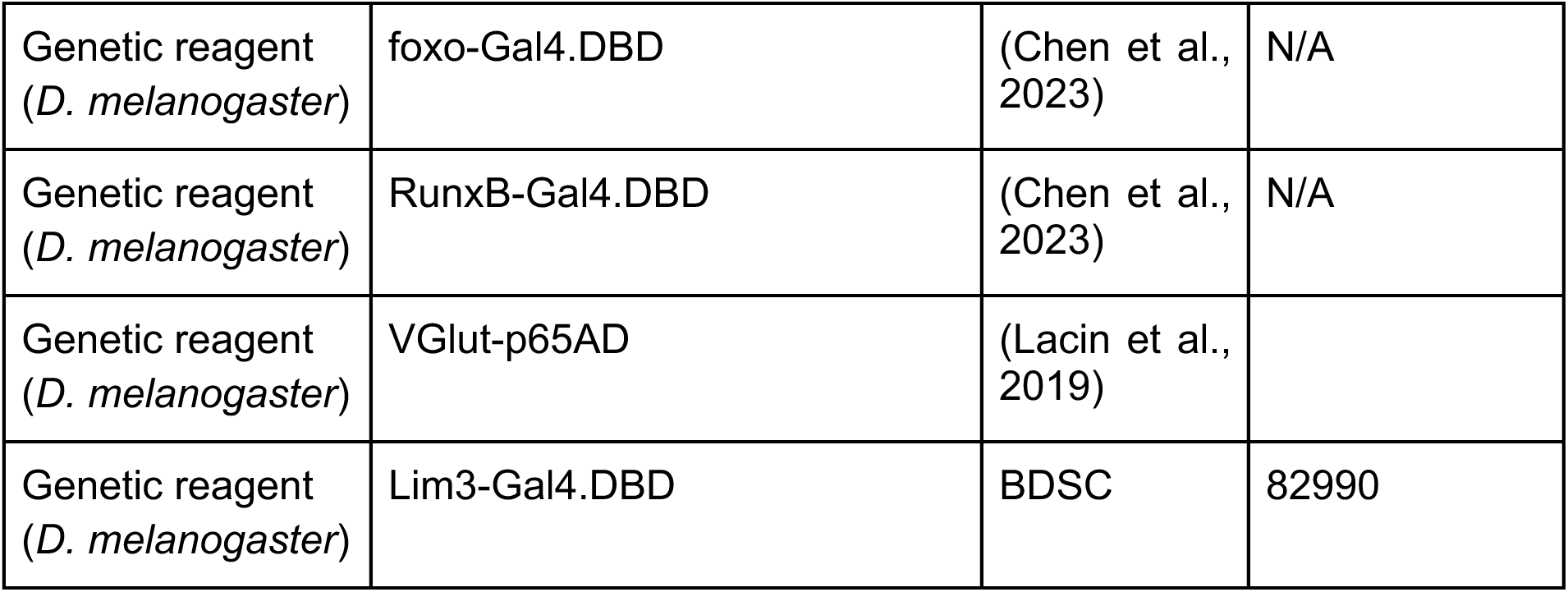

### Tools used for MN labeling

For scRNAseq, we genetically labeled all LinA-born neuroblast and their descendant cells using a LinA-specific R10c12-Gal4 driver crossed with the reporter immortalization tool (Awasaki et al., 2014). The progenies of this cross, including both male and female, were used for VNC dissections at four developmental time points: late L3 stage (∼96h after egg laying), 20 hours after puparium formation (APF), 45-hour APF and one-day-old adults. All the flies were maintained at 25°C.

### T2A-split-Gal4 method

CRISPR-mediated T2A-split-Gal4 knock-in for Zfh1-T2A-VP16 was performed by WellGenetics Inc. (Taipei, Taiwan). In brief, the gRNA sequence TTGGCAGGTGGGCGGCACTG[AGG] was cloned into U6 promoter plasmid(s). Cassette T2A-VP16AD-Zip-GMR-RFP, which contains T2A, Zip-VP16AD-Zip, SV40 3’UTR, a floxed GMR-RFP, and two homology arms were cloned into pUC57-Kan as donor template for repair. Targeting gRNAs and hs-Cas9 were supplied in DNA plasmids, together with donor plasmid for microinjection into embryos of control strain *w1118*. F1 flies carrying selection marker of GMR-RFP were further validated by genomic PCR and sequencing.

### Adult VNC and leg dissection and mounting

For the dissection of adult VNC and leg, we followed the protocol as described previously (Enriquez et al., 2018; Guan et al., 2022).

### Image acquisition of adult VNC and legs

Multiple 1µm-thick sections in the z-axis for adult VNC and legs were imaged using the Leica TCS SP5 II or Zeiss LSM 800 Confocal Microscope. All image processing and visualization were carried out using FIJI image processing software (Schneider et al., 2012).

### Single-cell RNA-sequencing of FACS-sorted cells

#### Sample preparation, single-cell RNA sequencing and clustering

Fluorescently labeled LinA-born MNs and glial cells were isolated from VNC dissected at four developmental time points (late L3 stage, 20 hours APF, 45 hours APF and one-day-old adults) using the protocol described previously (Hempel et al., 2007). Genetically labeled cells were FACS-sorted from other neuronal populations, and MNs and glial cells were separated based on size. Single-cell transcriptome profiling of MNs was performed using the method described in Macosko et al. (Macosko et al., 2015). For clustering, we used the modified version of Levitin et al. as described previously (Levitin et al., 2019; Mizrak et al., 2020). We used UMAP for visualization.

#### Highly variable genes

For MN scRNAseq datasets, highly variable genes were identified using single-cell hierarchical Poisson factorization (scHPF) method as described in Levitin et al. (Levitin et al., 2019). For preMNs, we used FindVariableFeatures of Seurat, which was previously used Allen et al. (Allen et al., 2020) for adult VNC scRNAseq clustering.

#### Pseudo time trajectory analysis

Force-directed motor neurons trajectories were generated using the method described by Mizrak et al. (Mizrak et al., 2020). We used the optimal transport method to annotate MNs from the adult stage to the late larval stage (Schiebinger et al., 2019).

#### Matching transcriptional clusters to LinA-borned MNs

For adult scRNAseq cluster annotation, we imaged adult leg to assess reporter expression of 20xUAS-6xGFP driven by MiMIC or CRIMIC-based split-T2A-Gal4 or T2A-Gal4 lines for sparsely expressed genes (as detailed in the Key Resource Table and **Supplementary Table**). MN axon morphology of these crosses was visually compared with the stereotyped axon targeting and morphological features defined for individual LinA MNs by Baek and Mann et al. (Baek & Mann, 2009).

#### ConnectionMiner

We utilized a previously published connectome for the same Drosophila VNC region (Lesser et al., 2024), capturing synaptic connectivity between identified cell types (or hemilineages) of MNs and preMNs. This was provided as a directed adjacency matrix, indicating the presence or absence of synaptic connectivity from cell type i to cell type j.

To integrate single-cell gene expression with the known connectome, we developed a factorization model, called **ConnectionMiner**, that aims to learn two unknowns:

1. A probabilistic assignment matrix *P* ∈ [0, 1]^*n*×*N*^, whose entries *P*_(*c,t*)_ represent the probability that single cell c belongs to connectome type t. Rows of P sum to 1, so each single cell’s probabilities across all N types sum to unity. Certain single cells have partially restricted columns in P if prior knowledge indicates they belong to a known subset of cell types.
2. A gene-gene interaction matrix *B* ∈ ℝ^*g*×*g*^ that captures how gene expression patterns linearly predict connectivity strength. Conceptually, B encodes which gene pairs are likely to be involved in synaptic wiring.

The core assumption is that the connectome C (an *N* × *N* matrix) can be approximated by:

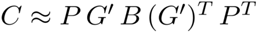

where *G′* ∈ ℝ^*n*×*g*^ is the single-cell expression matrix, and P maps from single-cell space to the N connectome-labeled types.

We iteratively update P and B to maximize the likelihood of reproducing C. The method enforces constraints on P:

- Rows sum to 1 (each single cell must “belong” fractionally to one or more cell types).
- Certain entries are fixed or restricted to zero if prior annotation excludes certain types.
- Additional regularization can be applied to B to encourage interpretability, such as sparsity and non-negativity. In this paper, we model B to be non-negative.

Upon convergence, each single cell c obtains a high probability in exactly one or a few cell types. This yields a refined or fully resolved assignment of single cells to connectome-defined types, thus bridging connectomic and transcriptomic data.

Once P is learned, we can compute the average expression profile *G* ∈ ℝ^*N*×*g*^ for each of the N connectome types:

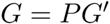

Hence, *G*_(*t,a*)_ is the average expression level of gene a in cell type t.

The schematic representation of ConnectionMiner factorization algorithm can be found in supplementary figure (Figure S4).

#### Validation of ConnectionMiner

To validate whether ConnectionMiner is learning the correct assignments of cells into cell types, we take a set of confidently annotated and known cell types (in this case, MNs Ti1, Ti9, and Fe5) and assume that their assignments are unknown. We then annotate all motor neurons into their cell types and then measure the correct assignment accuracy for the cell types that we are confident about. We observe that Ti1,Ti9 and Fe5 are assigned with about 74% confidence, where without connectivity information this rate is lower. This shows that ConnectionMiner is using the connectivity information to accurately assign single cells into their correct identities.

#### ConnectionMiner Interpretation

Once we ran ConnectionMiner, we could use the resultant G matrix and the connectome matrix C to perform univariate or bivariate statistical tests to identify gene pairs (*a, b*) strongly associated with synaptic connectivity. Specifically, for each pair of cell types (*t*_1_, *t*_2_), we label them as “connected” if *C*(*t*_1_, *t*_2_) = 1 and “not connected” otherwise. We compute expression features, e.g., coexpression = *G*(*t*_1_, *a*) × *G*(*t*_2_, *b*). Then we compare distributions of co-expression for connected vs. not-connected pairs using rank-sum or t-tests, generating a volcano plot of effect size vs. significance (-log p-values).

We further annotate genes into families (e.g., cell adhesion, receptor, synaptic, transcription factor, etc.). By grouping significant gene pairs by these categories, we compute average effect sizes or significance for each family, thus highlighting broad functional classes that drive connectivity.

To ensure computational efficiency given the large number of potential gene-gene pairs (∼5.38 million), our implementation uses a block-based and vectorized strategy to calculate statistics, along with optional parallel computing and optimal stopping criteria. Volcano plots we generate display each gene pair’s effect size (x-axis) vs. −log_10_(*p*) (y-axis). The p-values are adjusted for multiple comparisons by the Bonferroni method. We highlight the top pairs (indicated by their Pareto front) that combine large effect size and strong significance.

We produce a heatmap matrix whose rows represent pairs of cell types (*t*_1_ → *t*_2_) that are connected in the final connectome, and columns represent the top gene pairs. Each cell in the heatmap is colored by an expression-based score such as:

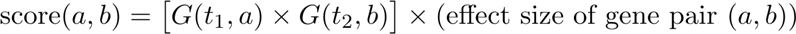

We highlight strong signals for particular connections and identify clusters of edges that share similar gene-pair usage.

Another visualization we used is a force-directed network diagram of preMNs and MNs. Each edge (*t*_1_ → *t*_2_) is expanded into K parallel edges, each corresponding to one of the top K gene pairs implicated in that connection. In our visualizations, we opt for K=3 to show top 3 gene pairs that have a high score for being implicated in a synapses (co-expression probability multiplied by effect size). We color these edges by the gene pair to visualize how multiple gene pairs might combine to support the same synapse. This helps guide future genetic knockdown or transTango experiments to disrupt specific gene pairs predicted to be crucial for particular edges.

#### Categorization of expressed and highly variable genes

We have categorized genes into different gene families: transcription factors (TFs), cell adhesion molecules (CAMs), ion channels (ICs), RNA binding genes (RBGs) and ribosomal proteins (RP) based on the definition described previously (ref). For other gene families, we used FlyBase ID, gene groups (GG) and gene ontology (GO) annotations obtained from FlyBase. Neuromuscular synaptic transmission (neuromuscular junction (NMJs); from GO:0007274). The chemoconnectome (CCT) gene family includes neurotransmitters, neuromodulators, neuropeptides (FBgg0000179), insulin-like peptides (FBgg0000048), transmembrane receptors (FBgg0001106) that include acetylcholine receptor, NMDA and non-NMDA ionotropic glutamate receptors, acetylcholine receptors, G protein-coupled receptors (neuroreceptors; FBgg0000172) excluding purine GPCRs, rhodopsins, unclassified class A GPCRs, class B GPCRs subfamily B2 and class F GPCRs. TFs gene families were further subclassified as basic helix-loop-helix TFs (bHLH; FBgg0000727), basic leucine zipper TFs (bZIPs; FBgg0000726), homeobox TFs (FBgg0000744) and zinc finger TFs (ZnF; FBgg0000729).

### Grouping of MNs

#### Motor modules

All 29 LinA-MNs have been categorized into 9 motor modules based on the FANC connectivity matrix (Lesser et al., 2024). These modules are coxa promotion (n=1; B1), femur reductor (n=1; Tr1), tibia flex A (n=8; Fe2, Fe4, Fe6, Fe7, Fe9, Fe11, Fe12, Ti11), tibia flex B (n=5; Fe8, Fe10, Fe13, Ti12, Ti13), tibia flex C (n=1; Ti10), long tendon muscles (ltm) DIP-alpha (n=2; Fe5, Ti9), ltm (n=9; Fe1, Fe3, Ti2, Ti3, Ti4, Ti5, Ti6, Ti7, Ti8), tarsus depressor retrodepressor (n=1; Ti14), and tarsus depressor DIP-alpha (n=1; Ti1).

#### Individual muscle

The 29 LinA-MNs can be classified into 8 distinct groups based on their innervation of specific muscles. These groups include: tergopleural promoter (n=1; B1), femur reductor (n=1; Tr1), long tendon muscle (ltm) 2 (n=3; Fe1, Fe3, Fe5), tibia flexor muscle (n=3; Fe2, Fe4, Fe6), accessory tibia flexor (n=7; Fe7 to Fe13), ltm 1 (n=8; Ti2 to Ti9), tarsus depressor muscle (n=5; Ti1, Ti11 to Ti14), and levator muscle (n=1; Ti10).

#### Muscle group

The 29 LinA-MNs can be categorized into 5 discrete clusters according to their innervation patterns with respect to specific muscle types. These groups are tergopleural promoter (n=1; B1), ltm (n=11; Fe1, Fe3, Fe5, Ti2 to Ti9), depressor muscle (n=8; Fe2, Fe4, Fe6, Ti1, Ti11 to Ti14), reductor muscle (n=8; Tr1, Fe7 to Fe13), and levator muscles (n=1; Ti10).

### Diversity index calculation

To quantify how effectively each gene family differentiates neuronal cell types, we compute a “Diversity Index” (DI) based on the multivariate differences in gene expression profiles. For each gene family of interest (e.g., TFs, CAMs, ion channels), a subset of highly variable genes is first identified. We then apply a diagonalization strategy to extract a set of genes whose expression most distinctly “diagonalizes” the cell-type expression matrix that is, the genes that are most enriched in one cell type relative to all others. We do this by performing a one-vs-all comparison of the cells x gene expression values assigned to one cell type vs. cells that belong to all others to determine the most enriched genes for that cell. We then retain the top K genes to generate a “diagonal” pattern heatmap.

Subsequently, for each gene family, the DI is calculated using a multivariate analysis of variance (MANOVA) framework. In practice, we compute the average pairwise Mahalanobis distance between the multivariate group means corresponding to different cell types. This measure serves as a robust index of how “diverse” the expression of a given gene family is across the neuronal subpopulations. A higher DI implies that the gene family in question contributes strongly to the differentiation of cell types and, by extension, may play a significant role in specifying synaptic connectivity patterns.

## Data and software availability

All raw and processed data of scRNAseq for MNs were uploaded to the GEO with accession number GSE290807. For adult VNC scRNAseq dataset, publicly accessible dataset GSE141807 was used. Annotated connectivity matrices for preMNs-preMNs and preMNs-LinA MNs were downloaded from publicly available datasets at GitHub (https://github.com/tuthill-lab/Lesser_Azevedo_2023). The source code for connectionMiner will be made available upon request or upon publication (whichever comes first).

## Acknowledgments

We thank Haluk Lacin and Benjamin White for generously sharing fly stocks; Mike Cleary and Cheng-Ting Chien for sharing antibodies; Aaron Allen and Stephen Goodwin for sharing their code for scRNAseq data analysis; members of the Mann lab and Varol lab for comments and suggestions. We are also very grateful to David Stern and the Quantitative Genomics Core at Janelia Research Campus where the single cell sequencing was done. This work was funded by NIH grants R01NS070644 and U19NS104655 to R.S.M.; R00MN128772 to E.V.; K99EY035757 to Y.C.D.C. and by an HHMI visitor project granted to David Stern and R.S.M. We thank John Tuthill, Wei-Chung Allen Lee and the FANC community for generating and proofreading the female adult nerve cord (NIH grant R01MH117808). A.W.A was supported by 1RF1NS128785-01 to John C. Tuthill.

## Supplementary Figures

**Figure S1.**
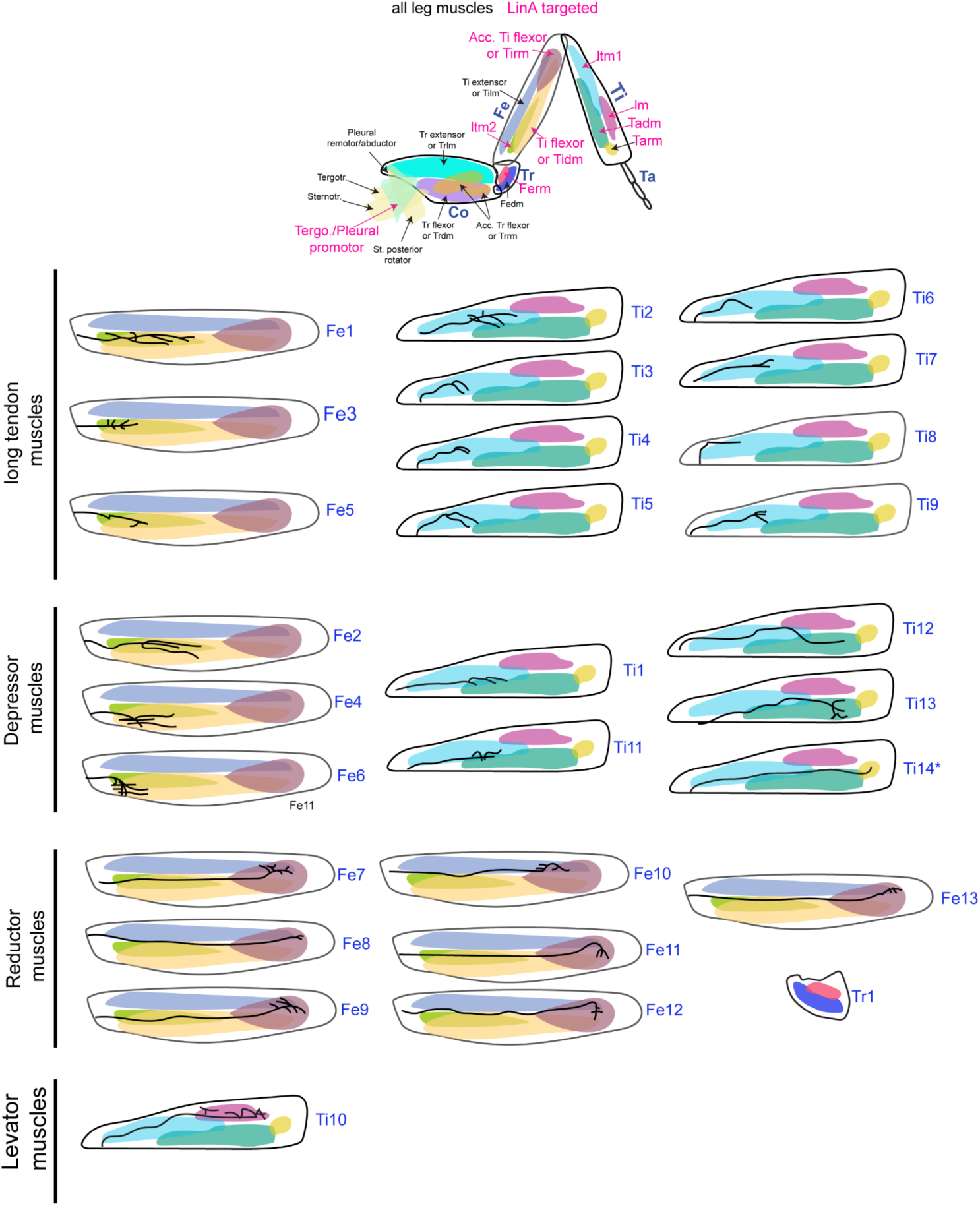
LinA generated leg MNs. Schematic showing stereotyped axon morphology and muscle targeting (highlighted as pink in top panel) of all LinA-born leg MNs in a front leg. MNs are arranged based on the muscle group they target. The schematic is modified from (Baek & Mann, 2009).

**Figure S2.**
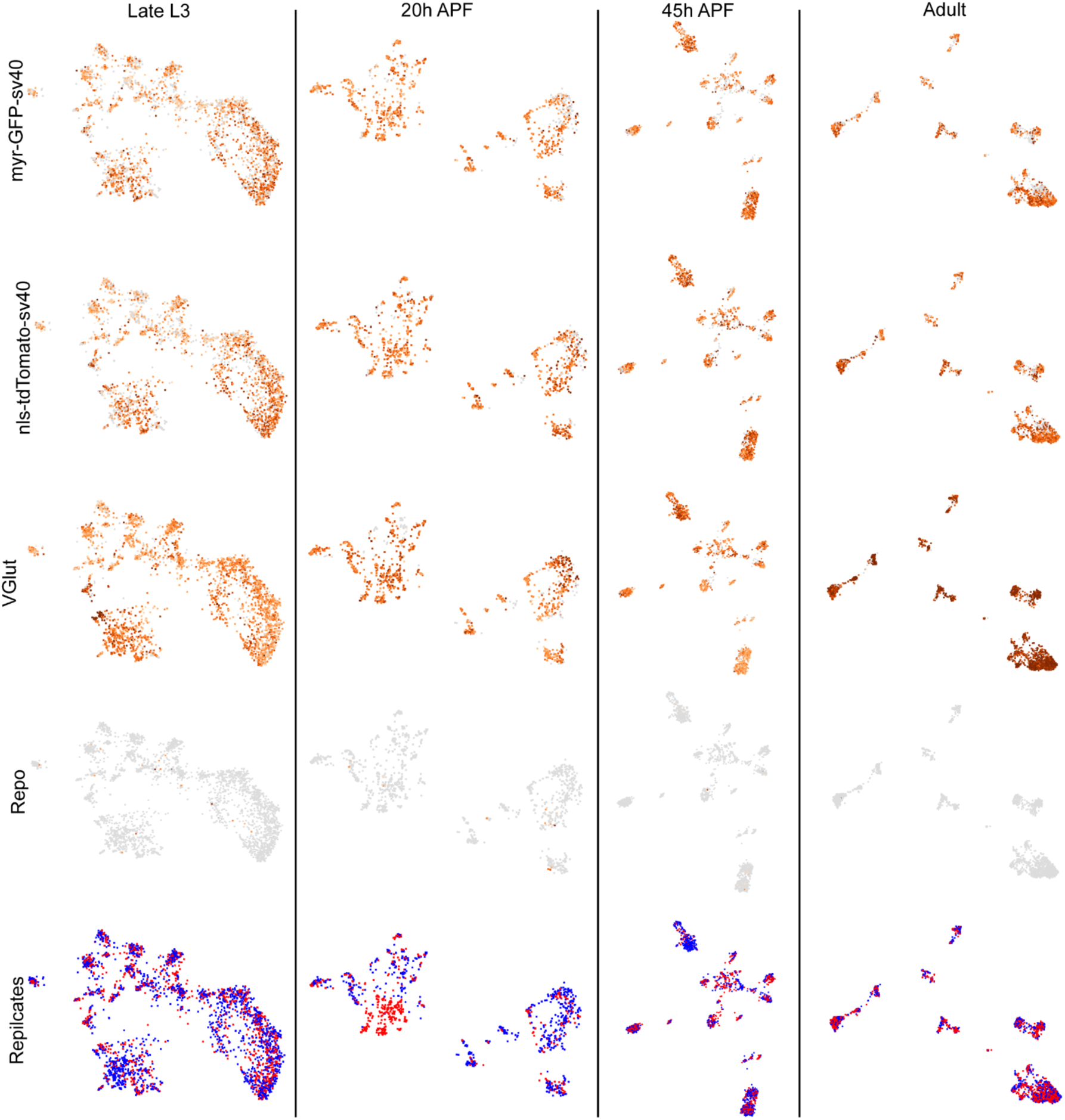
UMAP visualization. UMAP showing distribution of cells expressing genetic marker (myr-GFP-sv40 and nls-tdTomato), glutamergic marker (VGlut), glial marker (Repo) and representation of biological replicates at four time points - Late L3, 20hrs APF, 45hrs APF and one-day old adult flies.

**Figure S3.**
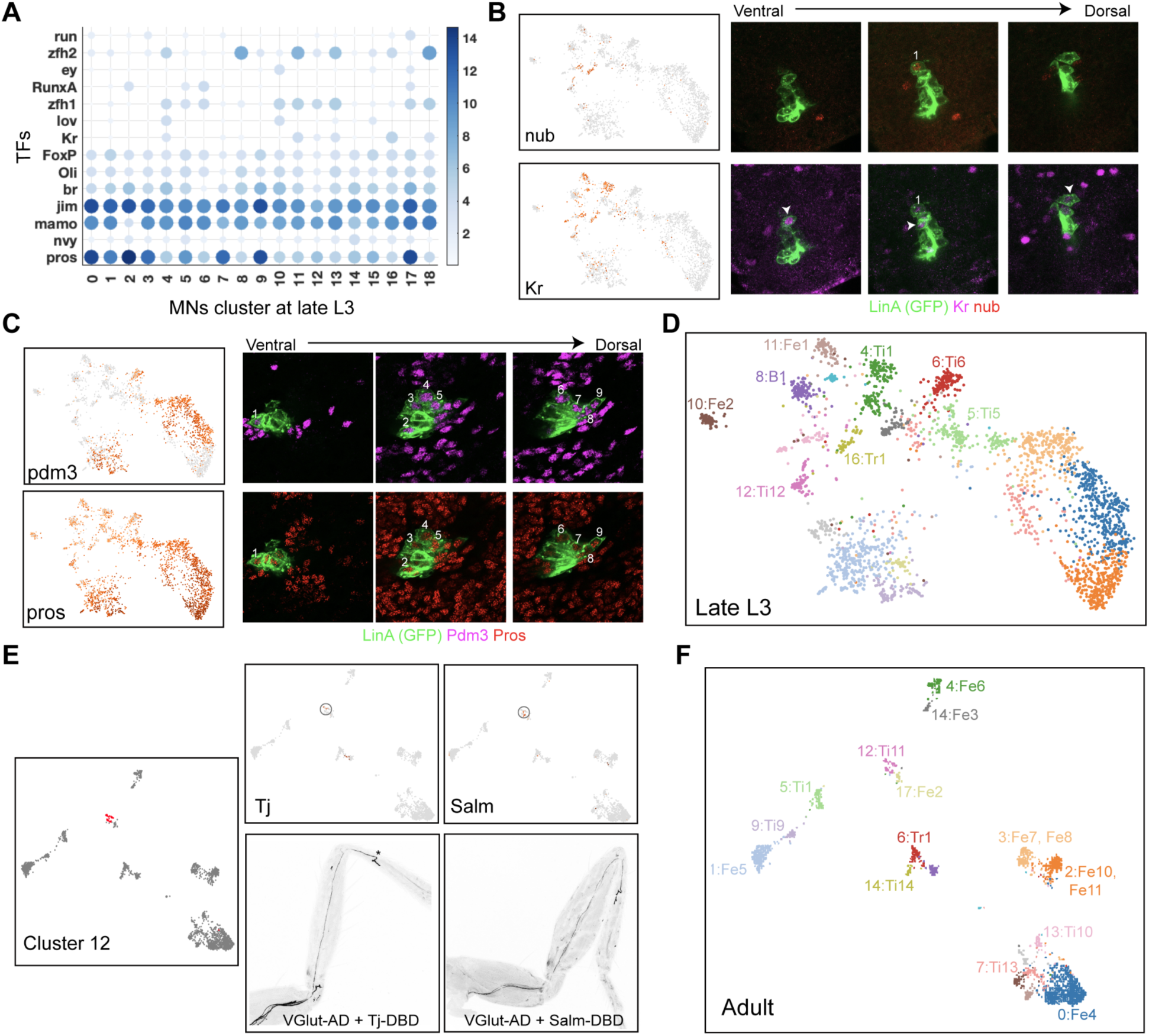
scRNAseq cluster annotation. (A) Heatmap showing expression of known morphological TFs (mTFs) in Late L3 stage scRNAseq clusters. (B) Immunostaining showing expression of nub protein (DGE in cluster 16) along with known mTF Kruppel (Kr) at the Late L3 stage. (C) Immunostaining showing expression of pdm3 protein (DGE in cluster 0, 1, 2, 3, 7, 9, 15 and 17) with known late-born MNs mTF marker prospero (pros) at Late L3 stage. (D) Annotation of 8 Late L3 scRNAseq cluster using immunostaining and mTFs expression. (E) UMAP showing cluster 12 in adult MNs scRNAseq dataset (left) and distribution of cells expressing Tj or Salm (upper panel). T2A-split-Gal4 for gene Tj or Salm, along with VGlut, shows their expression in Ti11 MN. (F) UMAP visualization of adult scRNAseq cluster showing partial annotation of 14 clusters before ConnectionMiner.

**Figure S4:**
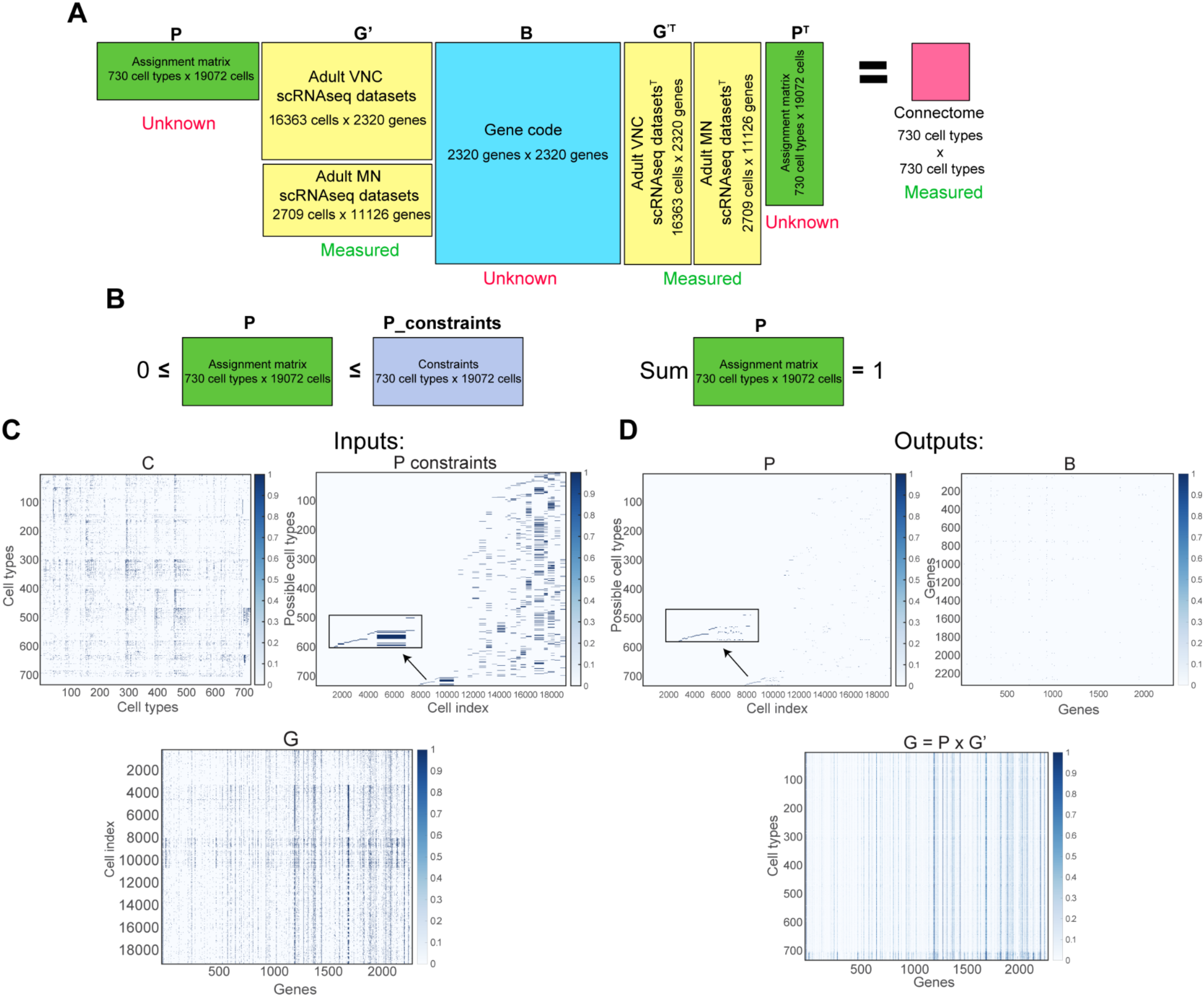
(A) Overall model. ConnectionMiner approximates the observed connectivity matrix “C” (rows and columns = cell types) as the product P × G′ × B × G′^T × P^T. Here “P” is an assignment matrix (rows = single cells, columns = cell types) that probabilistically maps each single cell to one or more cell types. “G′” is the single-cell gene-expression matrix, derived from ventral nerve cord (VNC) and motor neuron (MN) scRNAseq datasets. “B” is a gene-interaction matrix capturing how co-expression of particular genes influences the probability of synaptic connectivity. (B) ConnectionMiner constraints. Each factor in the model (“P” and “B”) is restricted to nonnegative values. “P” is additionally constrained by partial assignments from cluster or lineage annotations (plus morphological data), while “B” can be limited to known ligand–receptor or cell-surface molecule interactions. (C) Inputs. The method takes as input, a connectivity matrix “C” from an EM-based connectome, a gene-expression matrix “G′” from single-cell transcriptomics, and partial annotation constraints (e.g., known cell types, morphological markers). The insets illustrate sub-blocks of these matrices. (D) Outputs. ConnectionMiner jointly learns the assignment matrix “P,” refining each single cell’s type identity, and the gene-interaction matrix “B,” indicating which gene-gene co-expression events most strongly predict connectivity. Example heatmaps for “P” and “B” reveal newly assigned cell types and candidate wiring molecules that govern synaptic connections. Furthermore, we can obtain the cell type average gene expression by multiplying assignments P with single cell gene expression matrices: G = P x G’.

**Figure S5.**
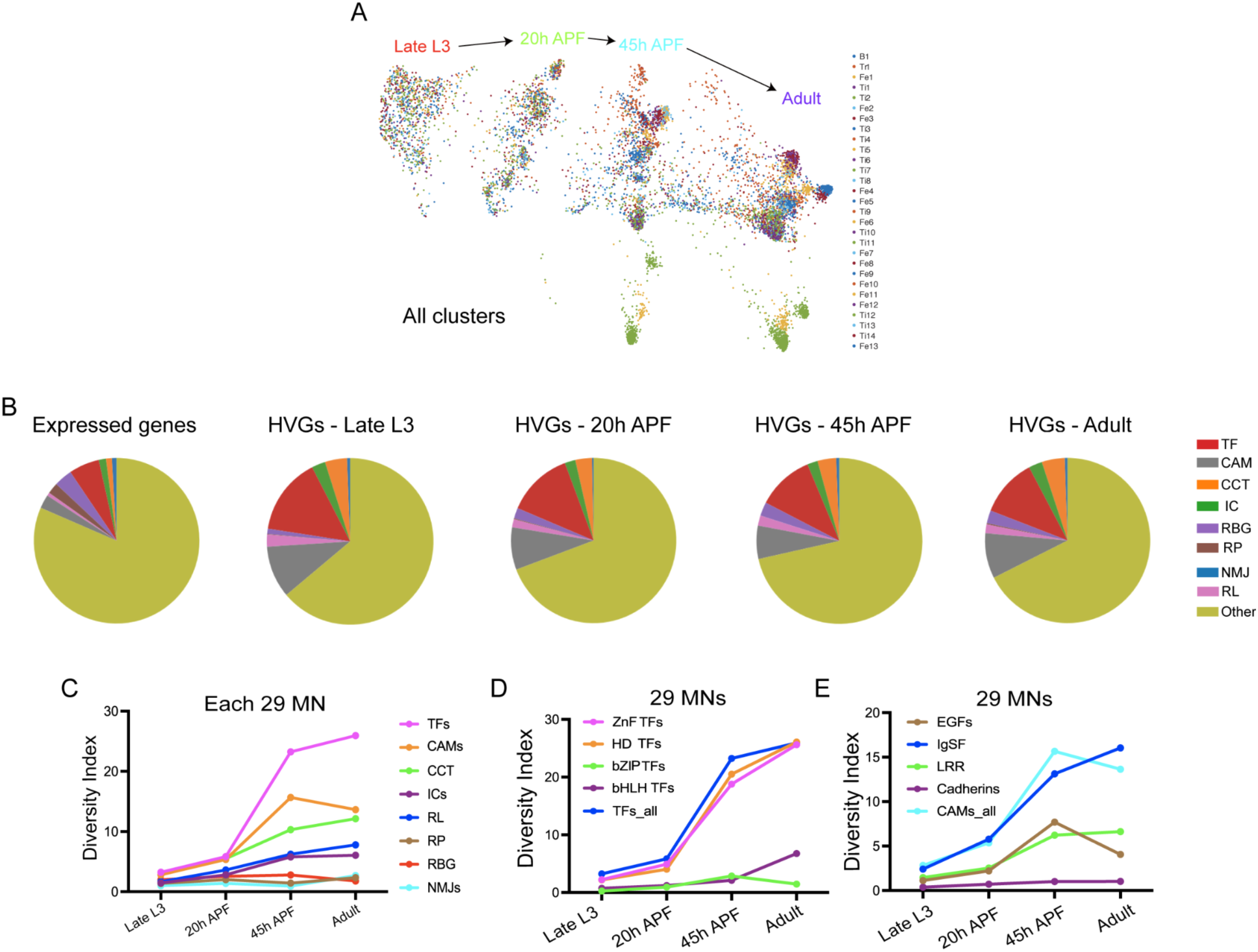
Optimal transport and gene distribution. (A) Optimal transport method showing distribution of cells belonging to same MN across four time points. (B) Distribution of expressed genes (left) and highly-variable genes (right) in different functional gene families in MNs at four developmental stages - Late L3, 20hrs APF, 45hrs APF and one-day old adult flies. Functional gene families are color coded.

**Figure S6.**
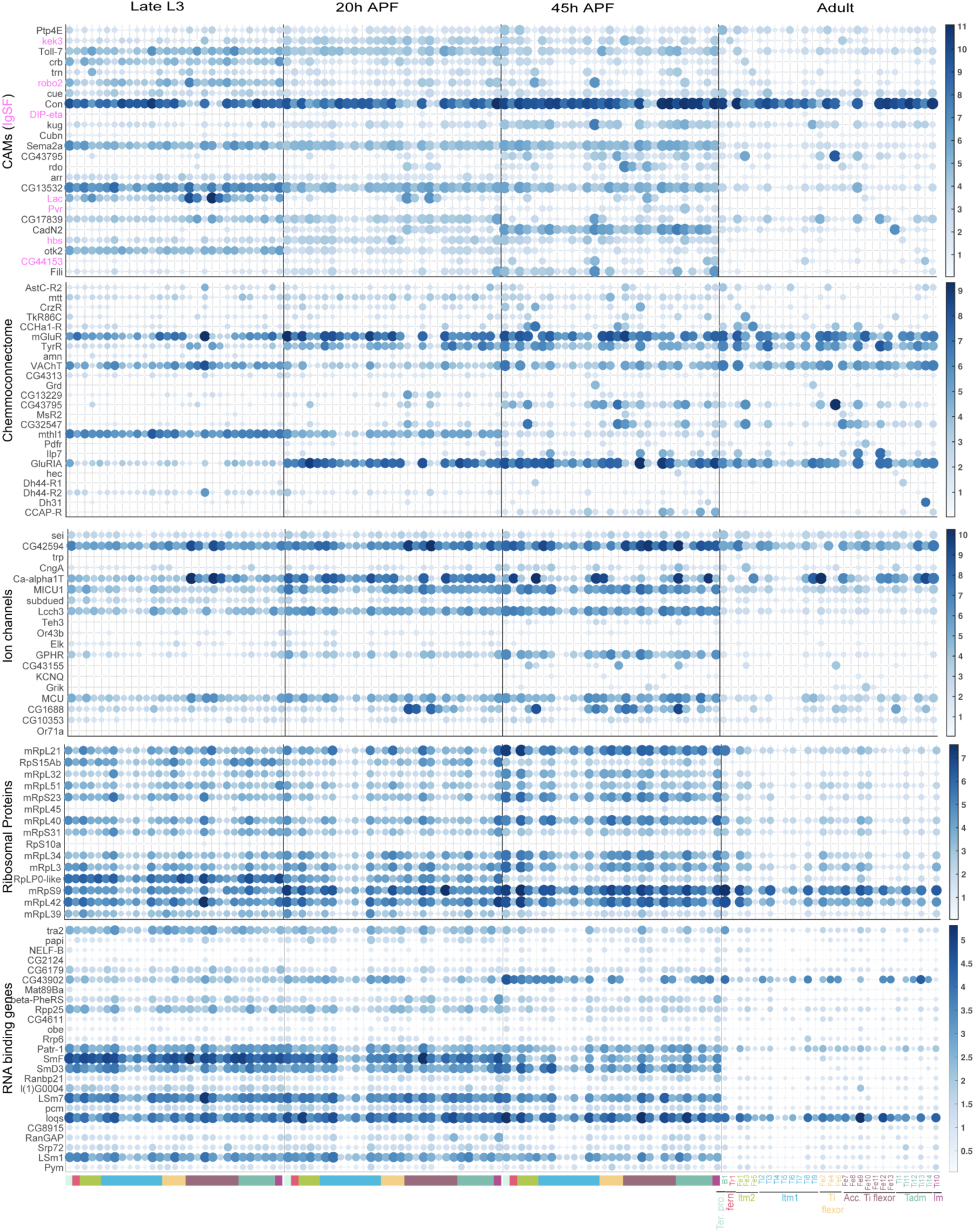
Top DEG for the 29 LinA-derived MNs. Heatmaps showing the expression of top 1 gene for different functional gene families in 29 leg MN in adults and their developmental expression patterns at other time points. MNs are arranged based on muscle innervation. MNs are color-coded based on the muscle they target.

**Figure S7.**
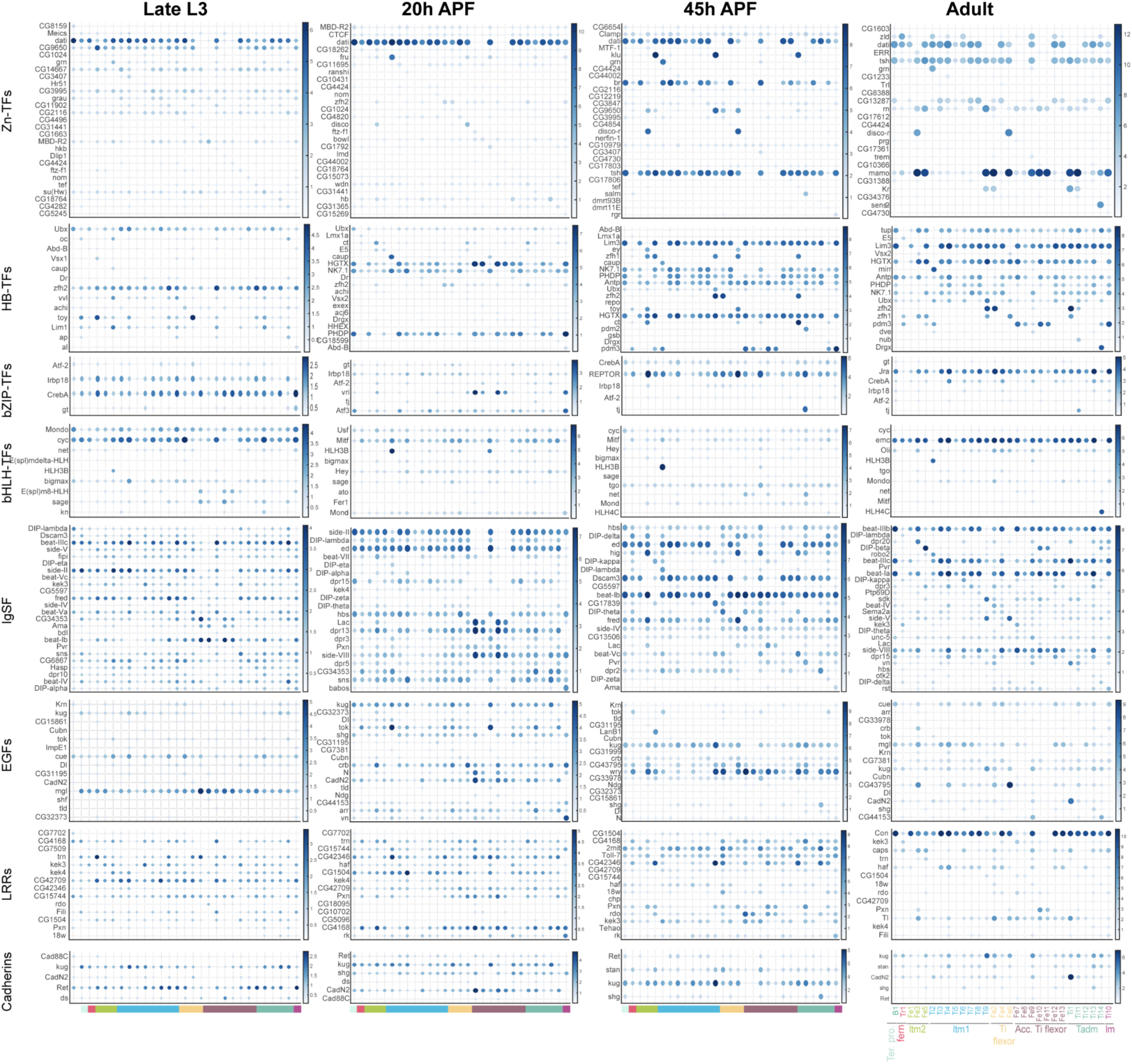
Top DEG in 29 MNs for TF and CAM gene subfamilies. Heatmaps showing the expression of top 1 gene for TFs and CAMs functional gene subfamilies in 29 leg MN in Late L3, 20hrs APF, 45hrs APF and adult time points.MNs are arranged based on muscle innervation. MNs are color-coded based on the muscle they target.

**Figure S8.**
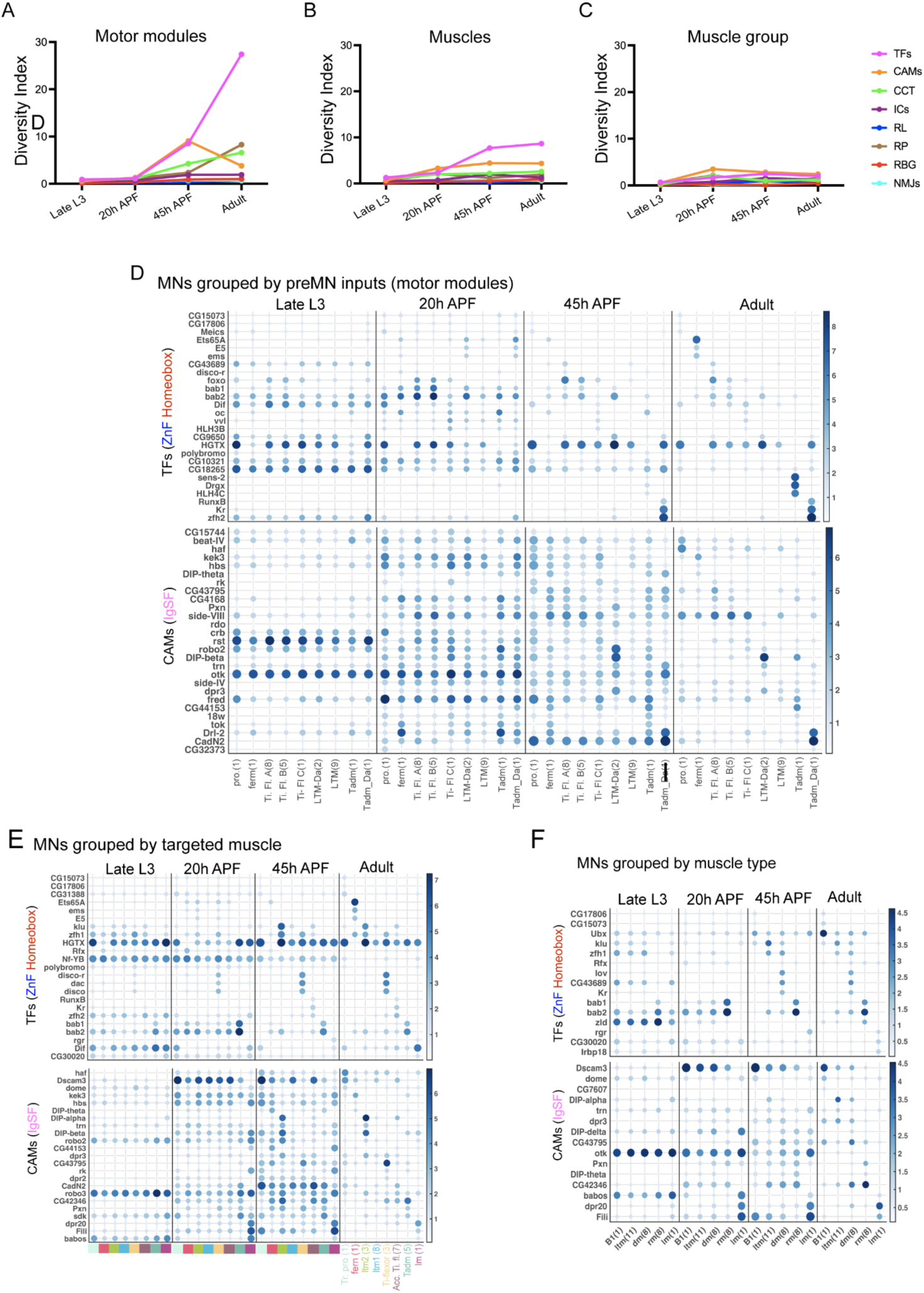
DEG for different groupings of MNs. (A-C) Graph showing diversity index values for different functional gene families at four developmental time points when MNs were grouped into motor modules (A), muscle (B) and muscle group (C). (D-F) Heatmap showing expression of top 3 TFs and CAMs in MNs when grouped into motor modules (D), muscle (E) and muscle group (F).

**Figure S9.**
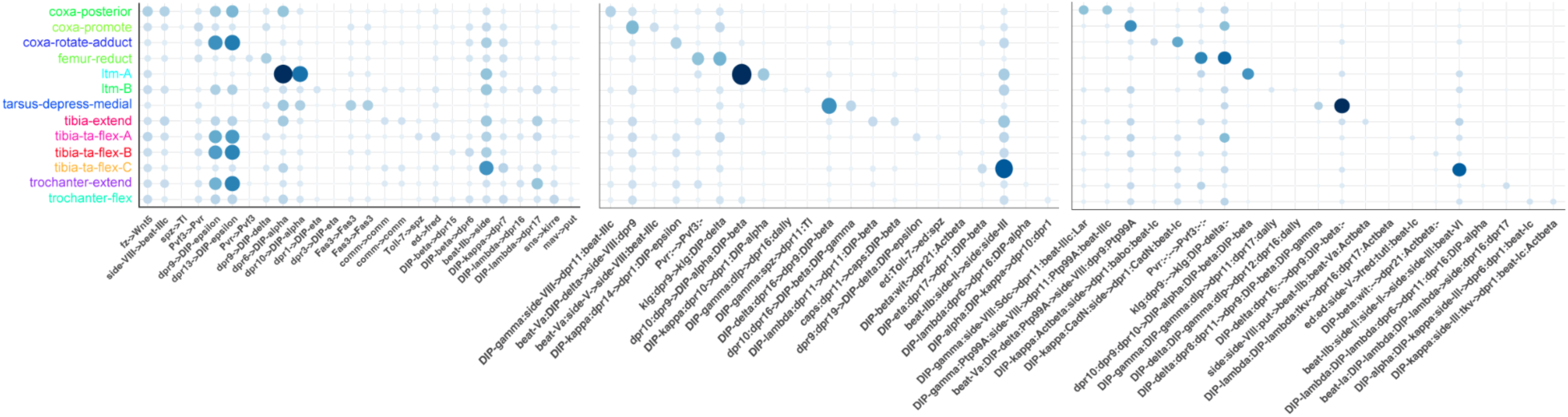
Columns indicate differentially expressed gene pairs in different premotor modules in the rows. The three plots show gene combinations that consider 1×1 gene pairs (left), as well as 2×2 (middle) and 3×3 (right), indicating the increasing levels of differential coding of premotor modules with higher order gene combinations.

## References

Allan, D. W., & Thor, S. (2015). Transcriptional selectors, masters, and combinatorial codes: Regulatory principles of neural subtype specification. WIREs Developmental Biology, 4(5), 505–528. 10.1002/wdev.191

Allen, A. M., Neville, M. C., Birtles, S., Croset, V., Treiber, C. D., Waddell, S., & Goodwin, S. F. (2020). A single-cell transcriptomic atlas of the adult Drosophila ventral nerve cord. eLife, 9, e54074. 10.7554/eLife.54074

Aran, D., Looney, A. P., Liu, L., Wu, E., Fong, V., Hsu, A., Chak, S., Naikawadi, R. P., Wolters, P. J., Abate, A. R., Butte, A. J., & Bhattacharya, M. (2019). Reference-based analysis of lung single-cell sequencing reveals a transitional profibrotic macrophage. Nature Immunology, 20(2), 163–172. 10.1038/s41590-018-0276-y

Awasaki, T., Kao, C.-F., Lee, Y.-J., Yang, C.-P., Huang, Y., Pfeiffer, B. D., Luan, H., Jing, X., Huang, Y.-F., He, Y., Schroeder, M. D., Kuzin, A., Brody, T., Zugates, C. T., Odenwald, W. F., & Lee, T. (2014). Making Drosophila lineage-restricted drivers via patterned recombination in neuroblasts. Nature Neuroscience, 17(4), 631–637. 10.1038/nn.3654

Azevedo, A., Lesser, E., Phelps, J. S., Mark, B., Elabbady, L., Kuroda, S., Sustar, A., Moussa, A., Khandelwal, A., Dallmann, C. J., Agrawal, S., Lee, S.-Y. J., Pratt, B., Cook, A., Skutt-Kakaria, K., Gerhard, S., Lu, R., Kemnitz, N., Lee, K., … Tuthill, J. C. (2024). Connectomic reconstruction of a female Drosophila ventral nerve cord. Nature, 1–9. 10.1038/s41586-024-07389-x

Azevedo, A. W., Dickinson, E. S., Gurung, P., Venkatasubramanian, L., Mann, R. S., & Tuthill, J. C. (2020). A size principle for recruitment of Drosophila leg motor neurons. eLife, 9, e56754. 10.7554/eLife.56754

Baek, M., & Mann, R. S. (2009). Lineage and birth date specify motor neuron targeting and dendritic architecture in adult Drosophila. The Journal of Neuroscience: The Official Journal of the Society for Neuroscience, 29(21), 6904–6916. 10.1523/JNEUROSCI.1585-09.2009

Barabási, D. L., & Barabási, A.-L. (2020). A Genetic Model of the Connectome. Neuron, 105(3), 435–445.e5. 10.1016/j.neuron.2019.10.031

Battistini, C., & Tamagnone, L. (2016). Transmembrane semaphorins, forward and reverse signaling: Have a look both ways. Cellular and Molecular Life Sciences: CMLS, 73(8), 1609–1622. 10.1007/s00018-016-2137-x

Bidaye, S. S., Machacek, C., Wu, Y., & Dickson, B. J. (2014). Neuronal Control of Drosophila Walking Direction. Science, 344(6179), 97–101. 10.1126/science.1249964

Brierley, D. j., Rathore, K., VijayRaghavan, K., & Williams, D. w. (2012). Developmental origins and architecture of Drosophila leg motoneurons. Journal of Comparative Neurology, 520(8), 1629–1649. 10.1002/cne.23003

Calleja, M., Moreno, E., Pelaz, S., & Morata, G. (1996). Visualization of Gene Expression in Living Adult Drosophila. Science, 274(5285), 252–255. 10.1126/science.274.5285.252

Card, G., & Dickinson, M. H. (2008). Visually Mediated Motor Planning in the Escape Response of Drosophila. Current Biology, 18(17), 1300–1307. 10.1016/j.cub.2008.07.094

Carrier, Y., Quintana Rio, L., Formicola, N., de Sousa-Xavier, V., Tabet, M., Chen, Y.-C. D., Ali, A. H., Wislez, M., Orts, L., Borst, A., & Pinto-Teixeira, F. (2024). Biased cell adhesion organizes the Drosophila visual motion integration circuit. Developmental Cell, S1534–5807(24)00638-5. 10.1016/j.devcel.2024.10.019

Carrillo, R. A., Özkan, E., Menon, K. P., Nagarkar-Jaiswal, S., Lee, P.-T., Jeon, M., Birnbaum, M. E., Bellen, H. J., Garcia, K. C., & Zinn, K. (2015). Control of Synaptic Connectivity by a Network of Drosophila IgSF Cell Surface Proteins. Cell, 163(7), 1770–1782. 10.1016/j.cell.2015.11.022

Chen, Y.-C. D., Chen, Y.-C., Rajesh, R., Shoji, N., Jacy, M., Lacin, H., Erclik, T., & Desplan, C. (2023). Using single-cell RNA sequencing to generate predictive cell-type-specific split-GAL4 reagents throughout development. Proceedings of the National Academy of Sciences of the United States of America, 120(32), e2307451120. 10.1073/pnas.2307451120

Cheong, H. S., Eichler, K., Stürner, T., Asinof, S. K., Champion, A. S., Marin, E. C., Oram, T. B., Sumathipala, M., Venkatasubramanian, L., Namiki, S., Siwanowicz, I., Costa, M., Berg, S., Janelia FlyEM Project Team, Jefferis, G. S., & Card, G. M. (2024). Transforming descending input into behavior: The organization of premotor circuits in the Drosophila Male Adult Nerve Cord connectome. 10.7554/eLife.96084.1

Clyne, J. D., & Miesenböck, G. (2008). Sex-Specific Control and Tuning of the Pattern Generator for Courtship Song in Drosophila. Cell, 133(2), 354–363. 10.1016/j.cell.2008.01.050

Cook, S. J., Jarrell, T. A., Brittin, C. A., Wang, Y., Bloniarz, A. E., Yakovlev, M. A., Nguyen, K. C. Q., Tang, L. T.-H., Bayer, E. A., Duerr, J. S., Bülow, H. E., Hobert, O., Hall, D. H., & Emmons, S. W. (2019). Whole-animal connectomes of both Caenorhabditis elegans sexes. Nature, 571(7763), 63–71. 10.1038/s41586-019-1352-7

Courgeon, M., & Desplan, C. (2019). Coordination between stochastic and deterministic specification in the Drosophila visual system. Science (New York, N.Y.), 366(6463), eaay6727. 10.1126/science.aay6727

Crickmore, M. A., & Vosshall, L. B. (2013). Opposing Dopaminergic and GABAergic Neurons Control the Duration and Persistence of Copulation in Drosophila. Cell, 155(4), 881–893. 10.1016/j.cell.2013.09.055

Dickinson, M. H., & Muijres, F. T. (2016). The aerodynamics and control of free flight manoeuvres in *Drosophila*. Philosophical Transactions of the Royal Society B: Biological Sciences, 371(1704), 20150388. 10.1098/rstb.2015.0388

Dombrovski, M., Zang, Y., Frighetto, G., Vaccari, A., Jang, H., Mirshahidi, P. S., Xie, F., Sanfilippo, P., Hina, B. W., Rehan, A., Hussein, R. H., Mirshahidi, P. S., Lee, C., Morris, A., Frye, M. A., Reyn, C. R. von, Kurmangaliyev, Y. Z., Card, G. M., & Zipursky, S. L. (2025). Gradients of Recognition Molecules Shape Synaptic Specificity of Visuomotor Transformation (p. 2024.09.04.610846). bioRxiv. 10.1101/2024.09.04.610846

Efremova, M., Vento-Tormo, M., Teichmann, S. A., & Vento-Tormo, R. (2020). CellPhoneDB: Inferring cell–cell communication from combined expression of multi-subunit ligand–receptor complexes. Nature Protocols, 15(4), 1484–1506. 10.1038/s41596-020-0292-x

Enriquez, J., Rio, L. Q., Blazeski, R., Bellemin, S., Godement, P., Mason, C., & Mann, R. S. (2018). Differing Strategies Despite Shared Lineages of Motor Neurons and Glia to Achieve Robust Development of an Adult Neuropil in Drosophila. Neuron, 97(3), 538–554.e5. 10.1016/j.neuron.2018.01.007

Enriquez, J., Venkatasubramanian, L., Baek, M., Peterson, M., Aghayeva, U., & Mann, R. S. (2015). Specification of individual adult motor neuron morphologies by combinatorial transcription factor codes. Neuron, 86(4), 955–970. 10.1016/j.neuron.2015.04.011

Guan, W., Bellemin, S., Bouchet, M., Venkatasubramanian, L., Guillermin, C., Laurençon, A., Kabir, C., Darnas, A., Godin, C., Urdy, S., Mann, R. S., & Enriquez, J. (2022). Post-transcriptional regulation of transcription factor codes in immature neurons drives neuronal diversity. Cell Reports, 39(13), 110992. 10.1016/j.celrep.2022.110992

Hebb, D. O. (2005). The Organization of Behavior: A Neuropsychological Theory. Psychology Press. 10.4324/9781410612403

Hempel, C. M., Sugino, K., & Nelson, S. B. (2007). A manual method for the purification of fluorescently labeled neurons from the mammalian brain. Nature Protocols, 2(11), 2924–2929. 10.1038/nprot.2007.416

>Hobert,oliver, & Kratsios, P. (2019). Neuronal identity control by terminal selectors in worms, flies, and chordates. Current Opinion in Neurobiology, 56. 10.1016/j.conb.2018.12.006

Hobert, O. (2016). Terminal Selectors of Neuronal Identity. Current Topics in Developmental Biology, 116. 10.1016/bs.ctdb.2015.12.007

Jakobsson, J. E. T., Spjuth, O., & Lagerström, M. C. (2021). scConnect: A method for exploratory analysis of cell–cell communication based on single-cell RNA-sequencing data. Bioinformatics, 37(20), 3501–3508. 10.1093/bioinformatics/btab245

Jin, S., Guerrero-Juarez, C. F., Zhang, L., Chang, I., Ramos, R., Kuan, C.-H., Myung, P., Plikus, M. V., & Nie, Q. (2021). Inference and analysis of cell-cell communication using CellChat. Nature Communications, 12(1), 1088. 10.1038/s41467-021-21246-9

Kovács, I. A., Barabási, D. L., & Barabási, A.-L. (2020). Uncovering the genetic blueprint of the C. elegans nervous system. Proceedings of the National Academy of Sciences, 117(52), 33570–33577. 10.1073/pnas.2009093117

Kurmangaliyev, Y. Z., Yoo, J., LoCascio, S. A., & Zipursky, S. L. (2019). Modular transcriptional programs separately define axon and dendrite connectivity. eLife, 8, e50822. 10.7554/eLife.50822

Kurmangaliyev, Y. Z., Yoo, J., Valdes-Aleman, J., Sanfilippo, P., & Zipursky, S. L. (2020). Transcriptional Programs of Circuit Assembly in the Drosophila Visual System. Neuron, 108(6), 1045–1057.e6. 10.1016/j.neuron.2020.10.006

Lacin, H., Chen, H.-M., Long, X., Singer, R. H., Lee, T., & Truman, J. W. (2019). Neurotransmitter identity is acquired in a lineage-restricted manner in the Drosophila CNS. eLife, 8, e43701. 10.7554/eLife.43701

Lacin, H., & Truman, J. W. (2016). Lineage mapping identifies molecular and architectural similarities between the larval and adult Drosophila central nervous system. eLife, 5, e13399. 10.7554/eLife.13399

Lee, P.-T., Zirin, J., Kanca, O., Lin, W.-W., Schulze, K. L., Li-Kroeger, D., Tao, R., Devereaux, C., Hu, Y., Chung, V., Fang, Y., He, Y., Pan, H., Ge, M., Zuo, Z., Housden, B. E., Mohr, S. E., Yamamoto, S., Levis, R. W., … Bellen, H. J. (2018). A gene-specific T2A-GAL4 library for Drosophila. eLife, 7, e35574. 10.7554/eLife.35574

Lesser, E., Azevedo, A. W., Phelps, J. S., Elabbady, L., Cook, A., Syed, D. S., Mark, B., Kuroda, S., Sustar, A., Moussa, A., Dallmann, C. J., Agrawal, S., Lee, S.-Y. J., Pratt, B., Skutt-Kakaria, K., Gerhard, S., Lu, R., Kemnitz, N., Lee, K., … Tuthill, J. C. (2024). Synaptic architecture of leg and wing premotor control networks in Drosophila. Nature, 1–9. 10.1038/s41586-024-07600-z

Levitin, H. M., Yuan, J., Cheng, Y. L., Ruiz, F. J., Bush, E. C., Bruce, J. N., Canoll, P., Iavarone, A., Lasorella, A., Blei, D. M., & Sims, P. A. (2019). De novo gene signature identification from single-cell RNA-seq with hierarchical Poisson factorization. Molecular Systems Biology, 15(2), e8557. 10.15252/msb.20188557

Li, H., Watson, A., Olechwier, A., Anaya, M., Sorooshyari, S. K., Harnett, D. P., Lee, H.-K. (Peter), Vielmetter, J., Fares, M. A., Garcia, K. C., Özkan, E., Labrador, J.-P., & Zinn, K. (2017). Deconstruction of the beaten Path-Sidestep interaction network provides insights into neuromuscular system development. eLife, 6, e28111. 10.7554/eLife.28111

Liu, J., Reggiani, J. D. S., Laboulaye, M. A., Pandey, S., Chen, B., Rubenstein, J. L. R., Krishnaswamy, A., & Sanes, J. R. (2018). Tbr1 instructs laminar patterning of retinal ganglion cell dendrites. Nature Neuroscience, 21(5), 659–670. 10.1038/s41593-018-0127-z

Lopez, D. H., Rostam, K., Zamurrad, S., Xu, S., & Mann, R. S. (2024). A critical affinity window for IgSF proteins DIP-α and Dpr10 is required for proper motor neuron arborization (p. 2024.09.11.612484). bioRxiv. 10.1101/2024.09.11.612484

Macosko, E. Z., Basu, A., Satija, R., Nemesh, J., Shekhar, K., Goldman, M., Tirosh, I., Bialas, A. R., Kamitaki, N., Martersteck, E. M., Trombetta, J. J., Weitz, D. A., Sanes, J. R., Shalek, A. K., Regev, A., & McCarroll, S. A. (2015). Highly Parallel Genome-wide Expression Profiling of Individual Cells Using Nanoliter Droplets. Cell, 161(5), 1202–1214. 10.1016/j.cell.2015.05.002

Menon, K. P., Kulkarni, V., Takemura, S.-Y., Anaya, M., & Zinn, K. (2019). Interactions between Dpr11 and DIP-γ control selection of amacrine neurons in Drosophila color vision circuits. eLife, 8, e48935. 10.7554/eLife.48935

Mizrak, D., Bayin, N. S., Yuan, J., Liu, Z., Suciu, R. M., Niphakis, M. J., Ngo, N., Lum, K. M., Cravatt, B. F., Joyner, A. L., & Sims, P. A. (2020). Single-Cell Profiling and SCOPE-Seq Reveal Lineage Dynamics of Adult Ventricular-Subventricular Zone Neurogenesis and NOTUM as a Key Regulator. Cell Reports, 31(12), 107805. 10.1016/j.celrep.2020.107805

Morano, N. C., Lopez, D. H., Meltzer, H., Sergeeva, A. P., Katsamba, P. S., Rostam, K. D., Gupta, H. P., Becker, J. E., Bornstein, B., Cosmanescu, F., Schuldiner, O., Honig, B., Mann, R. S., & Shapiro, L. (2024). Cis inhibition of co-expressed DIPs and Dprs shapes neural development. bioRxiv: The Preprint Server for Biology, 2024.03.04.583391. 10.1101/2024.03.04.583391

Morey, M., Sk, Y., T, H., A, N., E, B., & Sl, Z. (2008). Coordinate control of synaptic-layer specificity and rhodopsins in photoreceptor neurons. Nature, 456(7223). 10.1038/nature07419

Namiki, S., Ros, I. G., Morrow, C., Rowell, W. J., Card, G. M., Korff, W., & Dickinson, M. H. (2022). A population of descending neurons that regulates the flight motor of Drosophila. Current Biology, 32(5), 1189–1196.e6. 10.1016/j.cub.2022.01.008

Ohyama, T., Schneider-Mizell, C. M., Fetter, R. D., Aleman, J. V., Franconville, R., Rivera-Alba, M., Mensh, B. D., Branson, K. M., Simpson, J. H., Truman, J. W., Cardona, A., & Zlatic, M. (2015). A multilevel multimodal circuit enhances action selection in Drosophila. Nature, 520(7549), 633–639. 10.1038/nature14297

Özel, M. N., Simon, F., Jafari, S., Holguera, I., Chen, Y.-C., Benhra, N., El-Danaf, R. N., Kapuralin, K., Malin, J. A., Konstantinides, N., & Desplan, C. (2021). Neuronal diversity and convergence in a visual system developmental atlas. Nature, 589(7840), 88–95. 10.1038/s41586-020-2879-3

Özkan, E., Carrillo, R. A., Eastman, C. L., Weiszmann, R., Waghray, D., Johnson, K. G., Zinn, K., Celniker, S. E., & Garcia, K. C. (2013). An Extracellular Interactome of Immunoglobulin and LRR Proteins Reveals Receptor-Ligand Networks. Cell, 154(1), 228–239. 10.1016/j.cell.2013.06.006

Pavlou, H. J., Lin, A. C., Neville, M. C., Nojima, T., Diao, F., Chen, B. E., White, B. H., & Goodwin, S. F. (2016). Neural circuitry coordinating male copulation. eLife, 5, e20713. 10.7554/eLife.20713

Sanes, J. R., & Zipursky, S. L. (2020). Synaptic Specificity, Recognition Molecules, and Assembly of Neural Circuits. Cell, 181(3), 536–556. 10.1016/j.cell.2020.04.008

Scheffer, L. K., Xu, C. S., Januszewski, M., Lu, Z., Takemura, S., Hayworth, K. J., Huang, G. B., Shinomiya, K., Maitlin-Shepard, J., Berg, S., Clements, J., Hubbard, P. M., Katz, W. T., Umayam, L., Zhao, T., Ackerman, D., Blakely, T., Bogovic, J., Dolafi, T., … Plaza, S. M. (2020). A connectome and analysis of the adult Drosophila central brain. eLife, 9, e57443. 10.7554/eLife.57443

Schiebinger, G., Shu, J., Tabaka, M., Cleary, B., Subramanian, V., Solomon, A., Gould, J., Liu, S., Lin, S., Berube, P., Lee, L., Chen, J., Brumbaugh, J., Rigollet, P., Hochedlinger, K., Jaenisch, R., Regev, A., & Lander, E. S. (2019). Optimal-Transport Analysis of Single-Cell Gene Expression Identifies Developmental Trajectories in Reprogramming. Cell, 176(6), 1517. 10.1016/j.cell.2019.02.026

Schneider, C. A., Rasband, W. S., & Eliceiri, K. W. (2012). NIH Image to ImageJ: 25 years of image analysis. Nature Methods, 9(7), 671–675. 10.1038/nmeth.2089

Seeds, A. M., Ravbar, P., Chung, P., Hampel, S., Midgley, F. M., Jr, Mensh, B. D., & Simpson, J. H. (2014). A suppression hierarchy among competing motor programs drives sequential grooming in Drosophila. eLife, 3, e02951. 10.7554/eLife.02951

Sergeeva, A. P., Katsamba, P. S., Cosmanescu, F., Brewer, J. J., Ahlsen, G., Mannepalli, S., Shapiro, L., & Honig, B. (2020). DIP/Dpr interactions and the evolutionary design of specificity in protein families. Nature Communications, 11(1), 2125. 10.1038/s41467-020-15981-8

Soffers, J. H., Beck, E., Sytkowski, D. J., Maughan, M. E., Devasi, D., Zhu, Y., Wilson, B., Chen, Y.-C. D., Erclik, T., Truman, J. W., Skeath, J. B., & Lacin, H. (2025). A library of lineage-specific driver lines connects developing neuronal circuits to behavior in the Drosophila Ventral Nerve Cord (p. 2024.11.27.625713). bioRxiv. 10.1101/2024.11.27.625713

Stent, G. S. (1973). A Physiological Mechanism for Hebb’s Postulate of Learning. Proceedings of the National Academy of Sciences, 70(4), 997–1001. 10.1073/pnas.70.4.997

Sullivan, K. G., & Bashaw, G. J. (2023). Intracellular Trafficking Mechanisms that Regulate Repulsive Axon Guidance. Neuroscience, 508, 123–136. 10.1016/j.neuroscience.2022.07.012

Takemura, S., Hayworth, K. J., Huang, G. B., Januszewski, M., Lu, Z., Marin, E. C., Preibisch, S., Xu, C. S., Bogovic, J., Champion, A. S., Cheong, H. S., Costa, M., Eichler, K., Katz, W., Knecht, C., Li, F., Morris, B. J., Ordish, C., Rivlin, P. K., … Berg, S. (2023). *A Connectome of the Male* Drosophila *Ventral Nerve Cord*. 10.1101/2023.06.05.543757

Taylor, S. R., Santpere, G., Weinreb, A., Barrett, A., Reilly, M. B., Xu, C., Varol, E., Oikonomou, P., Glenwinkel, L., McWhirter, R., Poff, A., Basavaraju, M., Rafi, I., Yemini, E., Cook, S. J., Abrams, A., Vidal, B., Cros, C., Tavazoie, S., … Miller, D. M. (2021). Molecular topography of an entire nervous system. Cell, 184(16), 4329–4347.e23. 10.1016/j.cell.2021.06.023

Tuthill, J. C., & Wilson, R. I. (2016). Mechanosensation and adaptive motor control in insects. Current Biology : CB, 26(20), R1022. 10.1016/j.cub.2016.06.070

Venkatasubramanian, L., Guo, Z., Xu, S., Tan, L., Xiao, Q., Nagarkar-Jaiswal, S., & Mann, R. S. (2019). Stereotyped terminal axon branching of leg motor neurons mediated by IgSF proteins DIP-α and Dpr10. eLife, 8, e42692. 10.7554/eLife.42692

Winding, M., Pedigo, B. D., Barnes, C. L., Patsolic, H. G., Park, Y., Kazimiers, T., Fushiki, A., Andrade, I. V., Khandelwal, A., Valdes-Aleman, J., Li, F., Randel, N., Barsotti, E., Correia, A., Fetter, R. D., Hartenstein, V., Priebe, C. E., Vogelstein, J. T., Cardona, A., & Zlatic, M. (2023). The connectome of an insect brain. Science. 10.1126/science.add9330

Xu, C. S., Januszewski, M., Lu, Z., Takemura, S., Hayworth, K. J., Huang, G., Shinomiya, K., Maitin-Shepard, J., Ackerman, D., Berg, S., Blakely, T., Bogovic, J., Clements, J., Dolafi, T., Hubbard, P., Kainmueller, D., Katz, W., Kawase, T., Khairy, K. A., … Plaza, S. M. (2020). A Connectome of the Adult Drosophila Central Brain (p. 2020.01.21.911859). bioRxiv. 10.1101/2020.01.21.911859

Yazdani, U., & Terman, J. R. (2006). The semaphorins. Genome Biology, 7(3), 211. 10.1186/gb-2006-7-3-211

Yoo, J., Dombrovski, M., Mirshahidi, P., Nern, A., LoCascio, S. A., Zipursky, S. L., & Kurmangaliyev, Y. Z. (2023). Brain wiring determinants uncovered by integrating connectomes and transcriptomes. Current Biology, 33(18), 3998–4005.e6. 10.1016/j.cub.2023.08.020

Zhang, A. W., O’Flanagan, C., Chavez, E. A., Lim, J. L. P., Ceglia, N., McPherson, A., Wiens, M., Walters, P., Chan, T., Hewitson, B., Lai, D., Mottok, A., Sarkozy, C., Chong, L., Aoki, T., Wang, X., Weng, A. P., McAlpine, J. N., Aparicio, S., … Shah, S. P. (2019). Probabilistic cell-type assignment of single-cell RNA-seq for tumor microenvironment profiling. Nature Methods, 16(10), 1007–1015. 10.1038/s41592-019-0529-1

Zheng, Z., Lauritzen, J. S., Perlman, E., Robinson, C. G., Nichols, M., Milkie, D., Torrens, O., Price, J., Fisher, C. B., Sharifi, N., Calle-Schuler, S. A., Kmecova, L., Ali, I. J., Karsh, B., Trautman, E. T., Bogovic, J. A., Hanslovsky, P., Jefferis, G. S. X. E., Kazhdan, M., … Bock, D. D. (2018). A Complete Electron Microscopy Volume of the Brain of Adult Drosophila melanogaster. Cell, 174(3), 730–743.e22. 10.1016/j.cell.2018.06.019

